# Augmenting Radiation Sensitivity by Targeting PAR-Dependent Replication Fork Vulnerability in *IDH-*Mutant Glioma

**DOI:** 10.64898/2026.08.08.743634

**Authors:** Yosuke Kitagawa, Ali Nasser, Ami Kobayashi, Ethan Wetzel, Lisa Melamed, Chia-Chen Chang, Hiroaki Nagashima, Julie J. Miller, Hiroaki Wakimoto, Daniel P. Cahill

**Author notes:** **Corresponding Authors**, Hiroaki Wakimoto^1,2^, and Daniel P. Cahill^1,2^.

## Abstract

Mutations in isocitrate dehydrogenase 1 (*IDH1*) drive the early stages of gliomagenesis while simultaneously imposing replication stress that creates targetable vulnerabilities. Using both *in vitro* and *in vivo* models, we show that inhibition of poly(ADP-ribose) glycohydrolase (PARG) induces a poly(ADP-ribose) (PAR)-dependent augmentation of radiosensitivity in *IDH1*-mutant glioma cells. Metabolic repletion of NAD^+^ fails to rescue this effect, indicating that the vulnerability cannot be explained solely by NAD^+^ depletion. Instead, PARG inhibition profoundly alters replication fork progression and S-phase kinetics in *IDH1*-mutant cells. Mechanistically, ionizing radiation preferentially activates replication fork-associated damage response proteins DNA-dependent protein kinase catalytic subunit (DNA-PKcs) and X-ray repair cross-complementing protein 1 (XRCC1) in *IDH1*-mutant cells, a response partially reversed by pharmacologic inhibition of mutant *IDH1*. Importantly, pharmacologic inhibition of DNA-PKcs with AZD7648 during irradiation disrupts fork-associated repair signaling and markedly enhances cytotoxicity in *IDH1*-mutant glioma models. Together, these findings identify a PAR-dependent replication fork vulnerability that can be therapeutically exploited to selectively enhance radiosensitivity in *IDH1*-mutant gliomas.

**Statement of significance:** *IDH*-mutant gliomas harbor intrinsic replication stress yet lack targeted radiosensitization strategies. We identify a PAR-dependent replication fork vulnerability in which disruption amplifies radiation cytotoxicity by deregulating S-phase fork signaling. Pharmacologic DNA-PKcs inhibition exploits this dependency, providing a genotype-selective approach to enhance radiotherapy in *IDH*-mutant glioma.

## Introduction

Diffuse gliomas harboring mutations in isocitrate dehydrogenase 1 or 2 (*IDH1/2*) represent a molecularly defined and clinically distinct subset of primary brain tumors, encompassing both astrocytoma and oligodendroglioma under the current WHO classification framework. Affecting predominantly adults under the age of 50, *IDH*-mutant gliomas account for the majority of WHO grade 2–3 astrocytomas and constitute the most frequently diagnosed malignant primary brain tumors in this age group (1,2). Current standard-of-care, comprising maximal safe surgical resection, fractionated radiotherapy, and alkylating chemotherapy, has yielded meaningful improvements in progression-free and overall survival across landmark randomized trials (3,4). Yet despite these advances, tumor recurrence remains nearly universal, and effective salvage options at relapse are critically limited. The biological mechanisms that govern treatment resistance and disease progression in *IDH*-mutant gliomas are incompletely understood, and the therapeutic potential embedded within the tumor’s distinctive molecular vulnerabilities has not been fully exploited. Improving the durability of initial treatment response therefore represents one of the most pressing and consequential challenges in *IDH*-mutant gliomas.

Poly(ADP-ribose) (PAR) is a structurally heterogeneous, reversible post-translational modification synthesized by poly(ADP-ribose) polymerases (PARPs), principally PARP1, through iterative, NAD^+^-consuming ADP-ribosylation of substrate proteins, and dismantled primarily by poly(ADP-ribose) glycohydrolase (PARG), whose hydrolytic activity liberates ADP-ribose monomers, recycles the nucleotide pool, and resets downstream signaling (5–7). PAR turnover also shapes how cells adapt to genotoxic and replication stress by tuning PAR-dependent signaling dynamics (8). Critically, because PAR chain elongation consumes one NAD^+^ molecule per ADP-ribose addition, PARP hyperactivation imposes a direct and stoichiometric metabolic cost, and the pace of PARG-mediated PAR catabolism determines how rapidly free NAD^+^ is restored (5). These interlocking features, genomic signaling, metabolic coupling, and enzymatic reversibility, converge to establish the PARP-to-PARG axis as a multi-dimensional therapeutic vulnerability situated at the interface of DNA damage response and cellular energy metabolism.

*IDH*-mutant gliomas are among cancer types in which this vulnerability is most therapeutically tractable. Neomorphic IDH1/2 mutations divert the canonical oxidative decarboxylation of isocitrate toward the aberrant NADPH-consuming reduction of α -ketoglutarate, producing the oncometabolite R-2-hydroxyglutarate (2-HG) at supraphysiological intracellular concentrations (9,10). A key metabolic liability in this context is reduced NAD^+^ reserve, which can render *IDH*-mutant glioma cells vulnerable to interventions that further limit NAD^+^ availability (11). Importantly, this sensitivity is intrinsic to the mutant IDH genotype itself, not contingent on secondary co - mutations, making it a broadly applicable metabolic liability across IDH-mutant glioma irrespective of grade or molecular subtype. Building directly on this metabolic framework, we previously demonstrated that pharmacologic PARG inhibition potentiates the cytotoxicity of the alkylating agent temozolomide (TMZ) in IDH-mutant tumor cells by sequestering NAD^+^ within hyperaccumulated PAR chains and collapsing metabolic homeostasis (12). In that mechanistic model, TMZ-induced PARP activation transiently depletes free NAD^+^ in IDH-mutant cells whose biosynthetic reserve is already constrained, and concurrent PARG inhibition arrests this depleted state by blocking PAR catabolism—freezing NAD^+^ in polymeric form and sustaining the collapse. Confirmatory NAD^+^ rescue experiments demonstrated that metabolic lethality, rather than amplified DNA strand-break signaling alone, constitutes the primary cytotoxic mechanism in this combination (12). These findings established PAR turnover as a genotype-selective actionable node in IDH-mutant glioma, in which cellular metabolic fragility can be weaponized by pharmacologically uncoupling PAR synthesis from its degradation. Critically, however, this prior work was confined to the alkylation-induced DNA damage context. It remains an open and clinically pressing question whether the PARP–PARG axis analogously modulates IDH-mutant cellular responses to ionizing radiation, mechanistically distinct genotoxic insults, which generates a divergent spectrum of DNA lesions and imposes distinct demands on replication-coupled repair.

The DNA damage response landscape of IDH-mutant glioma is demonstrably altered, but its molecular determinants remain incompletely resolved and have been shown highly context-dependent. Multiple independent studies have documented perturbations in radiation sensitivity and DNA damage checkpoint engagement in IDH-mutant models (13–16). Sulkowski et al. reported that 2-HG competitively suppresses homologous recombination (HR) capacity and confers PARP inhibitor sensitivity (17). However, this HR-deficiency model has not been uniformly reproducible; other studies have failed to detect a consistent HR-deficient signature, and a mechanistic reexamination by Schvartzman et al. demonstrated that IDH1/2 mutation instead induces heterochromatin-dependent slowing of DNA replication fork progression, elevating replication stress and generating DNA double-strand breaks without obligate loss of HR pathway competence (18). This heterochromatin-driven replication stress model is important because it accounts for the breadth and context-dependence of the observed DNA damage phenotypes. Taken together, these findings argue compellingly against a single, obligatory DNA repair defect as the unifying vulnerability of IDH-mutant glioma, and instead support a reconceptualization in which dysregulated S-phase genome maintenance and impaired replication stress surveillance constitute a mechanistic and therapeutic vulnerability generalizable in *IDH*-mutant gliomas.

In this context, PAR-dependent replication signaling represents a mechanistic node linking mutant *IDH*-associated replication stress to therapeutic response. PAR signaling plays multiple key roles in S-phase genome maintenance. During unperturbed replication, PARP1 monitors lagging-strand synthesis by sensing ssDNA gaps at unligated Okazaki fragments, initiating localized PARylation that must be rapidly resolved by PARG to permit replication to proceed with fidelity (19). PARG-mediated dePARylation of proliferating cell nuclear antigen (PCNA) is required to restore its productive interaction with flap endonuclease 1 (FEN1), which is indispensable for Okazaki fragment maturation (20). Complementing this, PARP1 auto-modification plays an orthogonal regulatory role: auto-PARylation actively tempers replication fork speed and promotes faithful Okazaki fragment processing by modulating PARP1’s own chromatin retention (21). PARP1 PARylates the replisome scaffold TIMELESS (TIM), marking it for proteasome-dependent turnover that limits replication stress accumulation and stabilizes stalled forks, a checkpoint mechanism whose disruption amplifies fork instability (22). In parallel, at unresected stalled replication forks, PARP1 and the DNA-dependent protein kinase catalytic subunit (DNA-PKcs) engage cooperatively to recruit X-ray repair cross-complementing protein 1 (XRCC1), establishing a fork-associated repair platform essential for protection and restart (23). Viewed in its entirety, this molecular circuitry positions the PAR synthesis-to-breakdown turnover cycle as an indispensable mediator of S-phase genome integrity and provides a mechanistic rationale for investigating how PARG-regulated replication signaling shapes the distinctive genome maintenance vulnerabilities of IDH-mutant glioma cells under therapeutic stress.

Here, we extended these findings by combining genetic and pharmacologic inhibition of PARG to delineate the role of PAR-dependent replication control in *IDH* mutant gliomas. We show that disrupting PAR turnover, via *PARG* knockout or pharmacologic PARG inhibition, amplifies the cytotoxic effects of ionizing radiation (IR) specifically in *IDH*-mutant glioma cells. In addition, enhanced radiosensitivity in PARG-inhibited *IDH*-mutant glioma cells was accompanied by radiation-induced upregulation of replication fork resolution factors, XRCC1 and DNA-PKcs, a phenotype partially reversible upon mutant *IDH* inhibition. Importantly, we identified that this intrinsic replication-associated vulnerability can be therapeutically exploited by combining radiotherapy with DNA-PKcs inhibition. In addition, *IDH*-mutant cells exhibit baseline alterations in S-phase progression critically dependent on PAR homeostasis, both *in vitro* and *in vivo*. Together these findings highlight a rational combination strategy to enhance radiotherapeutic efficacy by targeting PAR-dependent replication folk vulnerability.

## Results

### PARG inhibition augments the cytotoxicity of irradiation in IDH mutant glioma cells

PARG inhibition, when combined with the alkylating agent TMZ, has been shown to induce selective cytotoxicity in *IDH*-mutant cancer cells (12). However, in our previous studies, we observed significantly attenuated NAD+-dependent cytotoxicity with the bifunctional alkylating agents carmustine (BCNU) and lomustine (CCNU), which function as DNA crosslinkers, compared to the monoalkylating agents TMZ and procarbazine (12). These findings indicated that the DNA damage response (DDR) vulnerability in *IDH*-mutant cells may be differentially engaged depending on the type of genotoxic insult. Since PARG inhibition has been shown to enhance radiation-induced cytotoxicity in breast cancer cells (24), we hypothesized that this combination might similarly augment radiation response in *IDH-*mutant glioma cells. Therefore, we investigated whether PARG inhibition could selectively enhance radiation-induced cytotoxicity in the context of *IDH1* mutation.

To investigate the effect of *IDH1* mutation, we first tested the combination of the highly selective PARG inhibitor PDD00017273 (hereafter PDD) with irradiation in MGG18 cells, an *IDH*-wild-type glioblastoma line engineered with a tetracycline-inducible *IDH1*-R132H transgene (Tet+/Tet−) (**Fig. 1A**). PDD selectively inhibited sphere formation in Tet+ cells (expressing IDH1-R132H) compared with Tet− cells (no IDH1-R132H expression), and this effect was further enhanced by irradiation (**Fig. 1A**). To assess the generalizability of our findings across endogenous *IDH*-mutant tumor models, we evaluated *IDH1*-R132C-mutant fibrosarcoma HT1080 cells. Consistent with our earlier results, treatment with 5 μM PDD enhanced radiation-induced cytotoxicity, as demonstrated by both cell viability and clonogenic survival assays (**Fig. 1B; Supplementary Fig. S1A and B**). Notably, this sensitizing effect was observed within a defined treatment window, specifically at 5 μM PDD in combination with 1 Gy irradiation, suggesting a dose-dependent interaction between PDD and radiation (**Fig. 1B**). However, in several patient-derived IDH-mutant glioma sphere lines, including MGG152, MGG119, and TS603, PARG inhibition enhanced IR cytotoxicity even at lower concentrations of PDD, while exhibiting a modest dose-dependent monotherapy effect (**Fig. 1C**).

**Figure 1.**
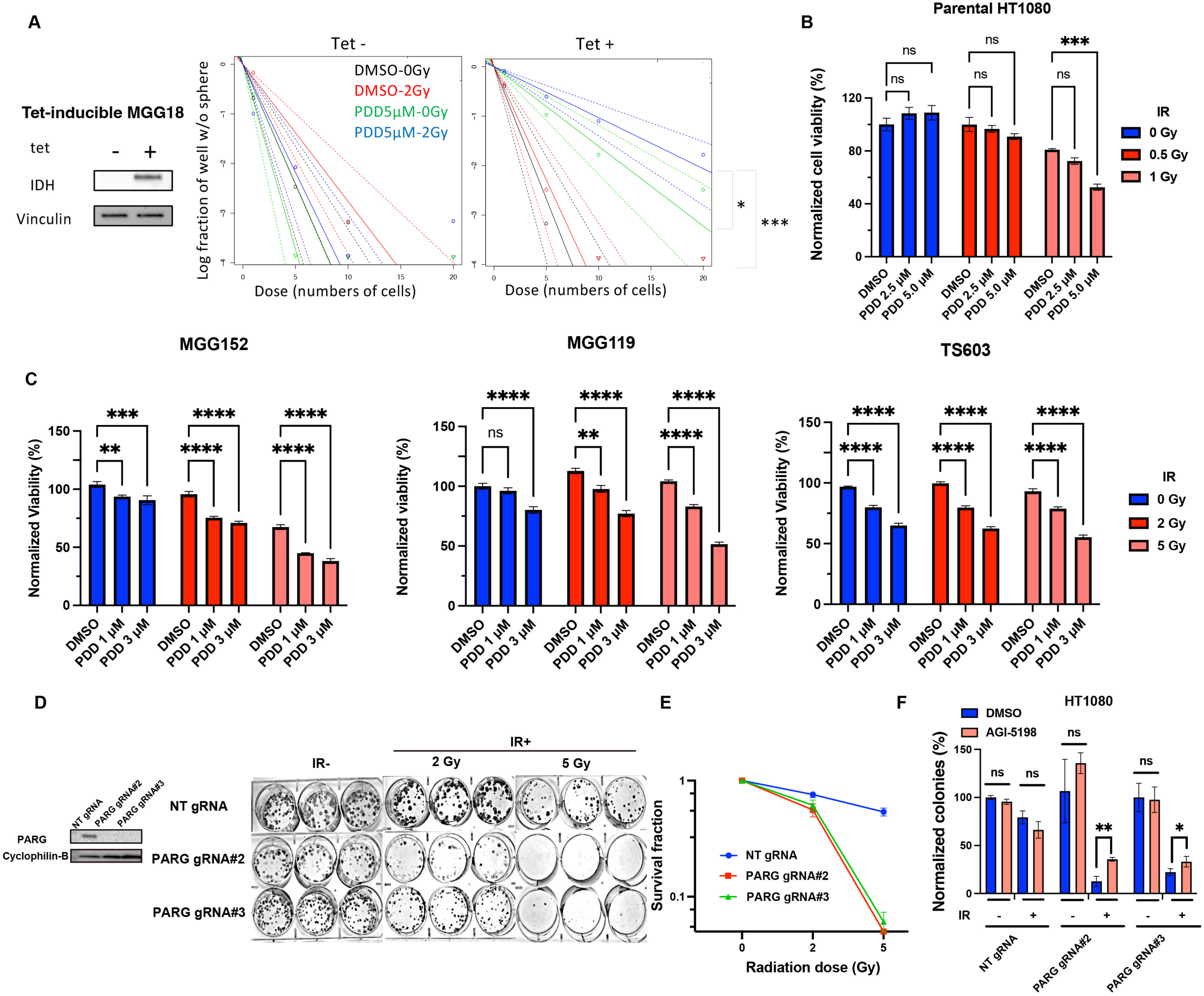

To validate on-target effects of PDD, we employed IDH1-mutant HT1080 cells with genetic ablation of PARG generated using two independent CRISPR guide RNAs (gRNA #2 and #3). Consistent with a direct role of PARG in modulating radiation response, PARG knockout (KO) cells exhibited a marked reduction in clonogenic survival following irradiation compared with non-targeting (NT) control cells (**Fig. 1D and E**). To further assess whether mutant IDH1 activity contributes to PARG loss–mediated radiosensitization, we treated PARG KO cells with the selective mutant IDH1 inhibitor AGI-5198. Notably, AGI-5198 partially rescued clonogenic survival in PARG KO HT1080 cells (**Fig. 1F; Supplementary Fig. S1C**), supporting a functional interaction between mutant IDH1 signaling and PARG-dependent responses to IR. Collectively, these results demonstrate that PARG inactivation enhances radiosensitivity in IDH-mutant tumor cells.

### DNA damage response, not NAD+ depletion, underlies the combination therapy with radiation and PARG inhibition in *IDH*-mutant cells

Given that *IDH-*mutant cells have reduced NAD^+^ reserves, and that DNA damage induced by IR activates PARP, leading to NAD^+^ consumption (11,12,24), we next investigated whether NAD^+^ metabolism contributes to the enhanced cytotoxicity observed with the combination of IR and PARG inhibition. To address this, we utilized β-nicotinamide mononucleotide (NMN), an intermediate in the NAD^+^ salvage biosynthesis pathway, to determine whether restoration of NAD^+^ levels could mitigate treatment-induced cytotoxicity (11). NMN supplementation, which increases intracellular NAD^+^ levels, significantly attenuated cytotoxicity in PARG KO HT1080 cells treated with TMZ, as well as in parental HT1080 cells treated with the PARG inhibitor PDD combined with TMZ (**Supplementary Fig. S2A-C**), consistent with prior reports (12). In contrast, NMN failed to rescue IR-induced suppression of colony formation in PARG KO HT1080 cells (**Fig. 2A and B**), suggesting that NAD^+^ depletion is not the primary driver of cytotoxicity in this context.

**Figure 2.**
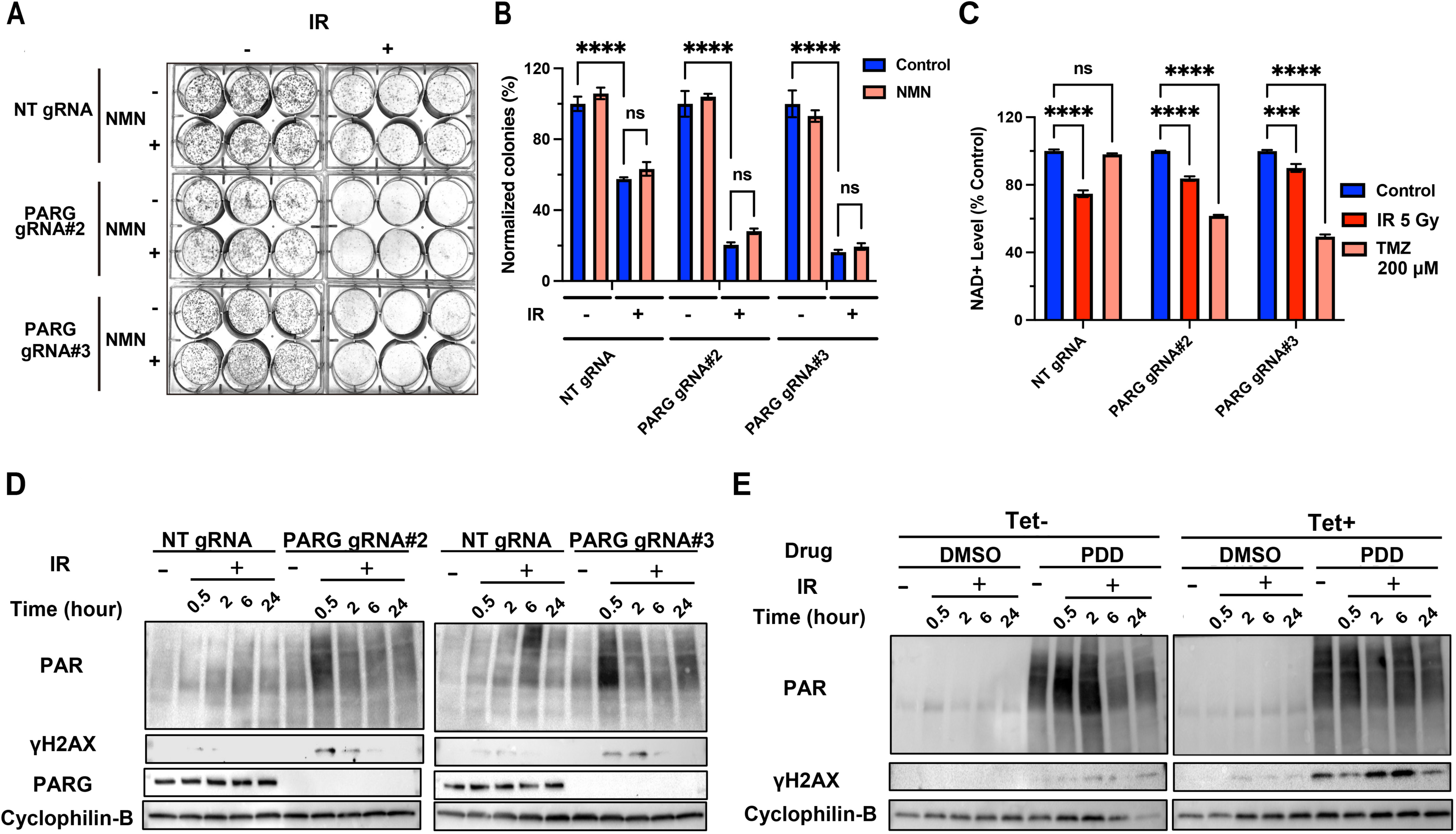

IR induced relatively modest reduction in NAD^+^ levels in both NT and PARG KO cells, whereas TMZ markedly decreased NAD^+^ levels specifically in PARG KO cells (**Fig. 2C**). These findings prompted us to investigate alternative mechanisms underlying the enhanced sensitivity to IR.

We therefore investigated the dynamics of PAR and γH2AX, markers of PARP activity and DNA damage, respectively. In PARG KO HT1080 cells, IR resulted in prolonged PAR accumulation and elevated γH2AX signals, indicating delayed signal resolution of DNA damage in the absence of PARG activity (**Fig. 2D**). Immunofluorescence (IF) analysis corroborated these findings. Both HT1080 NT and PARG KO cells exhibited increased PAR and γH2AX signals at 1 hour post-IR, displaying diffuse nuclear and cytoplasmic localization (**Supplementary Fig. S3A**), consistent with western blot of PAR (**Fig. 2D**). PAR and γH2AX signals were sustained and observed at 4 hours post-IR in PARG KO cells, but these declined in the presence of AGI-5198 (**Supplementary Fig. S3B and C**). In MGG18 cells, induction of IDH1 R132H expression (Tet^+^) did not alter PAR or γH2AX kinetics following IR, as responses were comparable to Tet⁻ cells (**Supplementary Fig. S3D-F**). However, upon combined treatment with PDD and IR, Tet^+^ cells exhibited prolonged PAR and γH2AX signals relative to Tet⁻ cells, and very modest γH2AX signals in response to IR and DMSO (**Fig. 2E**). Consistent with these observations, comet assays confirmed persistent DNA damage after IR in *IDH1*-mutant PARG KO cells, as evidenced by significantly elongated tail moments (**Supplementary Fig. S4A and B**). Together, these results indicate that the enhanced sensitivity to IR in the context of PARG loss is not primarily driven by NAD^+^ depletion. Instead, PARG deficiency leads to sustained PARylation and prolonged DNA damage signaling in response to IR in an IDH-mutant selective manner. These findings support a model in which aberrant PAR metabolism, rather than NAD^+^ depletion per se, underlies the persistent DDR observed in IDH-mutant cells following IR.

### PARG loss and mutant *IDH1* inhibition alter replication fork dynamics, S-phase progression, and replication-associated damage responses

Recent studies have suggested that replication fork stalling is more prevalent in *IDH*-mutant cells due to heterochromatin alterations (18). To directly assess replication fork progression, we performed sequential chlorodeoxyuridine (CldU)–iododeoxyuridine (IdU) pulse-labeling DNA fiber assays in NT and PARG-deficient HT1080 cells **(Fig. 3A)**. This dual-labeling approach enables discrimination of ongoing forks (CldU→IdU tracks) from stalled forks or newly fired origins, while total tract length (CldU + IdU) serves as a measure of folk progression. Under basal conditions, PARG knockout cells exhibited significantly increased total replication tract length compared with NT cells, indicating enhanced fork progression (**Fig. 3B and C**). Treatment with the mutant IDH1 inhibitor AGI-5198 further increased replication tract length in PARG KO cells beyond the elongation observed with PAR*G* loss alone (**Fig. 3B and C**). These findings suggest that both PARG deficiency and pharmacologic inhibition of mutant IDH1 activity promote replication fork progression.

**Figure 3.**
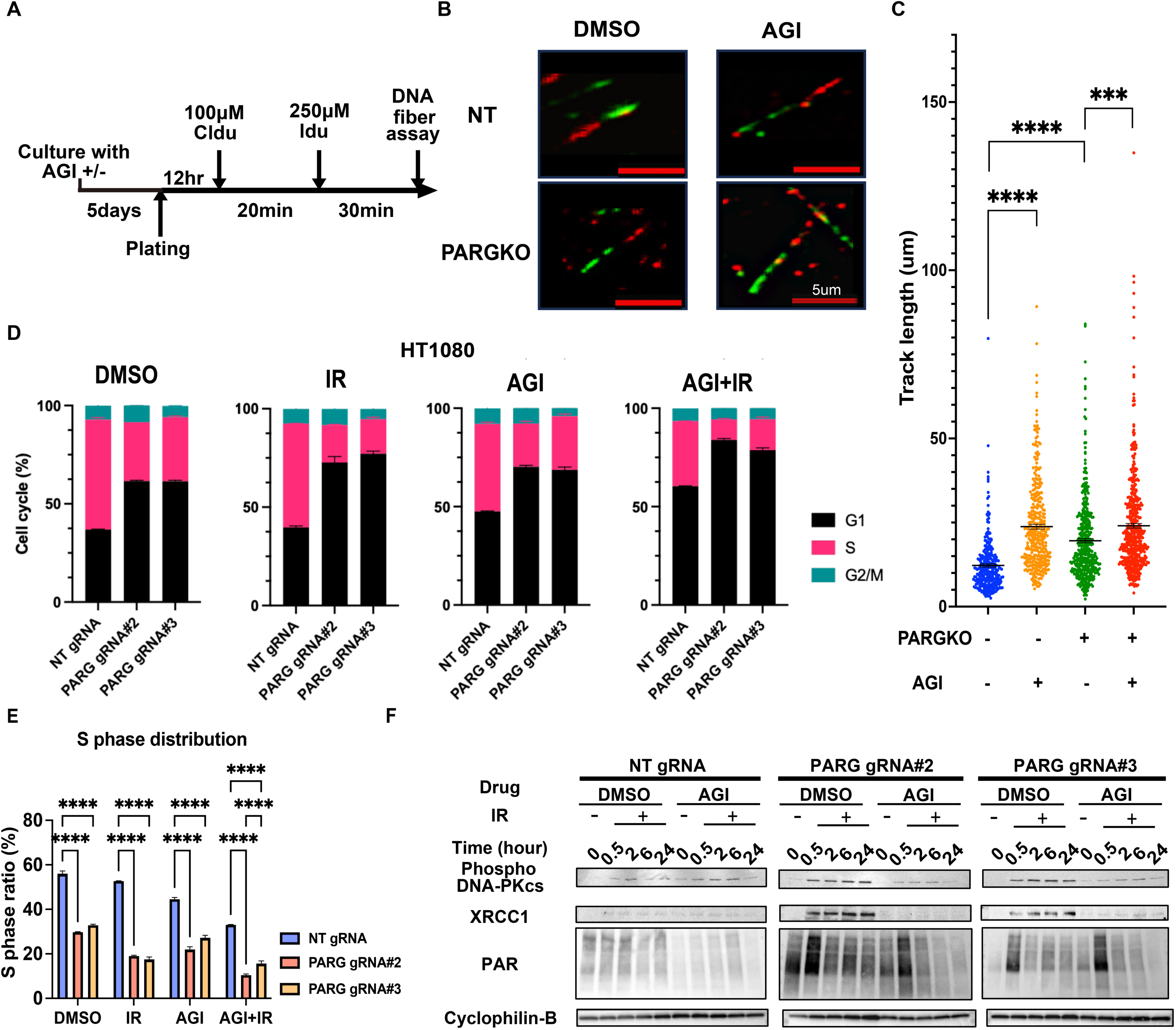

5-Ethynyl-2ʹ-deoxyuridine (EdU)–based cell-cycle analysis was performed using a predefined single-cell gating strategy, and results were quantified as S-phase fraction (see Methods). Comparison of cell-cycle profiles revealed pronounced shift in G1/S distribution in PARG KO cells relative to NT controls, consistent with an altered S-phase distribution (**Fig. 3D and E**). Upon IR exposure, cell-cycle profiling revealed further shift toward G1/S accumulation in PARG KO cells (**Fig. 3D and E**). Five days of AGI-5198 treatment reduced the S-phase fraction more prominently in NT cells than in PARG KO cells **(Fig. 3D and E)**. These data indicate that PARG loss promotes replication progression and decreases the S-phase fraction, which is further altered by IR.

Based on these alterations in S-phase dynamics, we next investigated replication fork–associated DDR signaling. In PARG KO cells, IR induced robust and sustained PAR accumulation, accompanied by marked induction of XRCC1 and phosphorylated DNA-PKcs as early as 0.5 h post-IR, persisting up to 24 h **(Fig. 3F)**. In contrast, under AGI-5198 treatment, IR-induced PAR accumulation was transient, and XRCC1 and phospho–DNA-PKcs signals were markedly reduced in PARG KO cells. In NT cells, these DDRs were comparatively modest and largely unaffected by AGI-5198 (**Fig. 3F**).

In parental HT1080 cells, IR alone induced strong γH2AX and phospho–DNA-PKcs signals, confirming activation of the DDR (**Supplementary Fig. S5A and B**). Notably, co-treatment with AGI-5198 did not significantly alter the induction of these markers immediately post-IR (**Supplementary Fig. S5A and B**), indicating that AGI-5198 does not mitigate the initial co-activation of phospho-DNA-PKcs and γH2AX. Collectively, these results demonstrate that IR-sensitivity in PARG deficient, IDH-mutant tumor cells is mechanistically associated with aberrant PAR-driven, replication stress S phase signaling.

### Inhibition of DNA-PKcs selectively augments radiation cytotoxicity in *IDH*-mutant cancer cells

Given the involvement of PARP1, XRCC1, and DNA-PKcs in stabilizing stalled replication forks in response to DNA damage (23), we next tested whether pharmacologic inhibition of DNA-PKcs could enhance radiation cytotoxicity, as observed with PARG inhibition. AZD7648 is a potent and highly selective DNA-PKcs inhibitor previously shown to enhance radiation cytotoxicity in breast cancer cells (25), although a genetic basis for tumor-selective radiosensitization has not been described. We investigated the combination of AZD7648 and IR in HT1080 NT and PARG KO cells. In addition to the expected enhanced sensitivity to IR in PARG KO cells, the combination of AZD7648 and IR produced a robust cytotoxic effect particularly in PARG KO cells (**Fig. 4A**). AZD7648 alone (up to 0.5 μM) had no effect on cell growth in our Tet-inducible MGG18 system, regardless of *IDH1*-R132H induction status (**Fig. 4B**). In contrast, the combination of AZD7648 and IR yielded clear *IDH*-specific radiosensitization, with the *IDH1*-R132H–expressing cells showing significantly reduced viability compared with *IDH1* wild-type cells (**Fig. 4B**). We extended these findings to patient-derived glioma sphere lines harboring endogenous *IDH1*-R132H mutations, including MGG152, MGG119, and TS603, and similarly observed that AZD7648 sensitized these cells to IR (**Fig. 4C–E**). For comparison, we tested the combination of AZD7648 with CCNU. However, the co-treatment with AZD7648 showed limited sensitization to CCNU in *IDH1*-mutant glioma sphere cells (**Supplementary Fig. S6A–C**). We attempted to reverse the effects of AZD7648 combination with IR using AGI-5198; however, co-incubation with AGI-5198 did not result in any discernible change in TS603 cell viability (**Supplementary Fig. S6D**).

**Figure 4.**
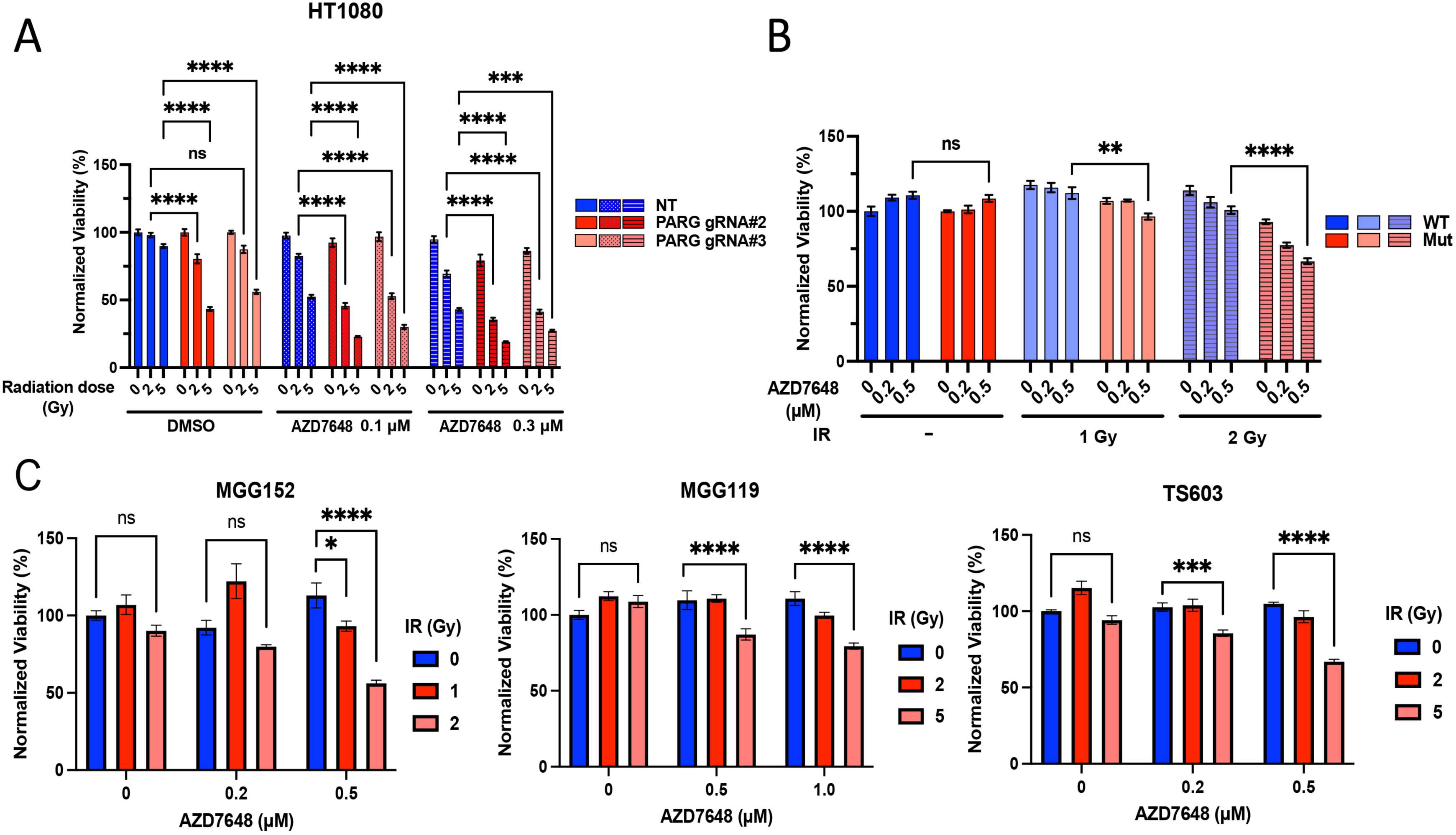

### PARG inhibition enhances radiosensitivity in an *IDH*-mutant cancer model *in vivo*

To further investigate the therapeutic effects of combining PARG loss with IR *in vivo*, we employed a bilateral flank xenograft model established with HT1080 cells expressing either NT gRNA or PARG KO (PARG-/-) in which only one tumor site was irradiated and the other site non-irradiated (**Fig. 5A; Supplementary Fig. S7A**). Once tumors reached approximately 100 mm³, one flank tumor per mouse received focal 3 Gy IR, while the contralateral tumor was left non-irradiated, and tumor growth and body weight were monitored longitudinally. In NT tumors, growth kinetics were comparable between irradiated and non-irradiated sites, indicating 3 Gy IR elicited minimal effect in this context (**Fig. 5B and C**). In contrast, in PARG KO tumors, which exhibited a slower growth rate compared with NT tumors, IR resulted in a marked suppression of tumor growth (**Fig. 5B and C**). Irradiated PARG KO tumors exhibited a significant reduction in tumor volume compared with their contralateral non-irradiated counterparts, demonstrating enhanced radiosensitivity upon PARG loss (**Fig. 5B and C**). Importantly, body weight remained stable across all groups throughout the 2.5-week post-irradiation period, indicating that observed anti-tumor effect was not associated with systemic toxicity (**Supplementary Fig. S7B**). To investigate the molecular basis of this differential response, we assessed DNA-damage and repair signaling markers in tumor tissues. IR induced elevated levels of PAR, XRCC1, phospho–DNA-PKcs, and γH2AX in PARG KO tumors (**Fig. 5D**), consistent with enhanced DNA-damage signaling and persistence of unrepaired lesions in the absence of PARG. Histopathological analysis by hematoxylin and eosin (H&E) staining revealed no overt morphological differences between NT and PARG KO tumors (**Fig. 5E**). However, quantitative analysis of Ki-67 immunostaining demonstrated a significant reduction in the proliferative index specifically in irradiated PARG KO tumors compared with all other groups (**Fig. 5E and F**). Together, these data demonstrate that PARG deficiency sensitizes tumors to IR *in vivo*, leading to enhanced tumor growth inhibition without detectable systemic toxicity. This effect is associated with sustained DNA damage signaling and reduced proliferative capacity, supporting a therapeutic strategy targeting PARG to potentiate radiotherapy responses.

**Figure 5.**
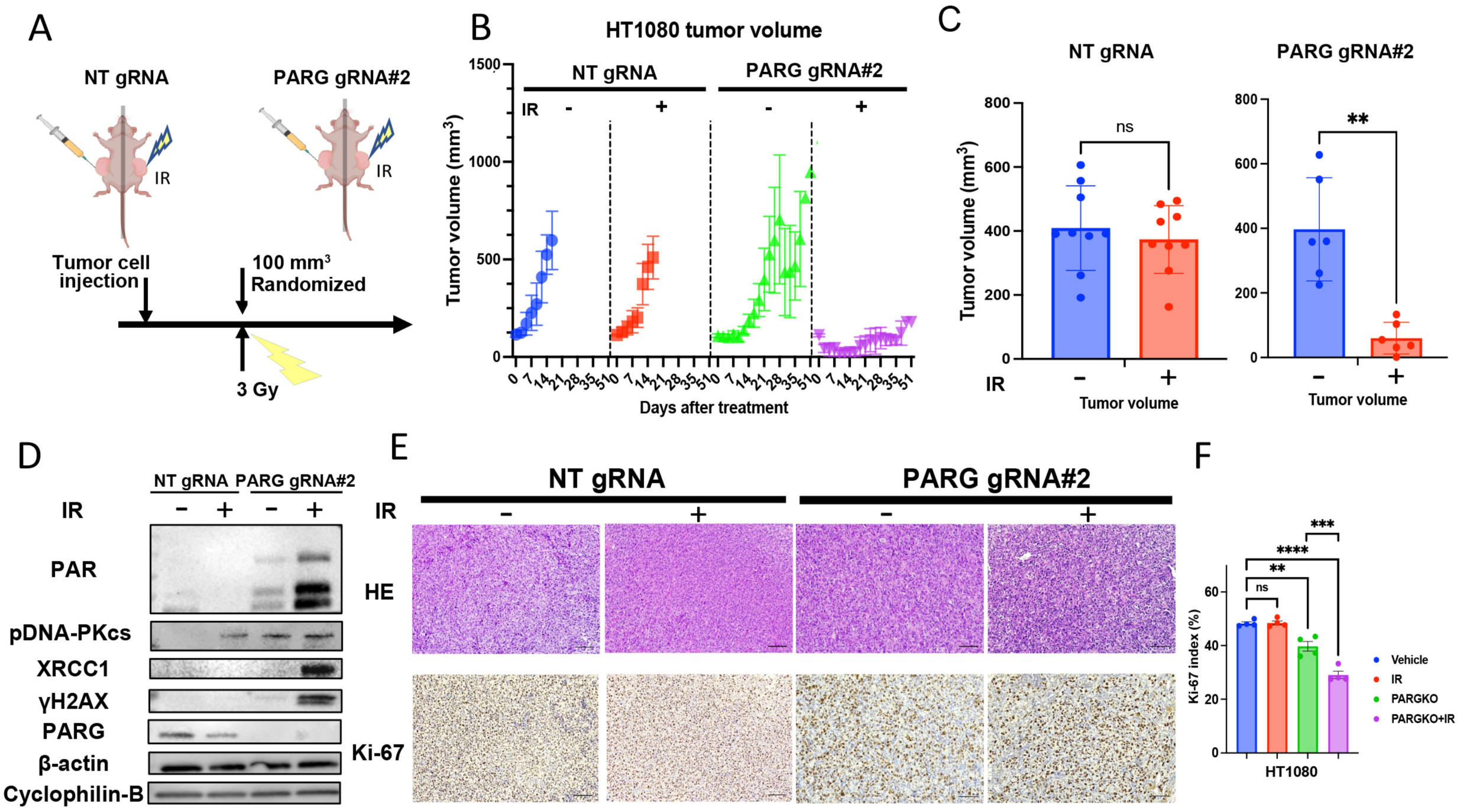

### DNA-PKcs inhibitor AZD7648 enhances radiosensitivity in an IDH-mutant cancer model *in vivo*

We next evaluated whether pharmacologic inhibition of DNA-PKcs using AZD7648 could sensitize *IDH1*-mutant tumors to IR *in vivo*, using the glioma-derived TS603 model. Tumors were established in a bilateral flank configuration to ensure equivalent systemic drug exposure in irradiated and non-irradiated sites within the same animal. Once tumors reached 80–120 mm³, mice were randomized to treatment groups and administered AZD7648 for five consecutive days, with IR administered on days 3 and 5 **(Fig. 6A, Supplementary Fig. S7C).** AZD7648 monotherapy did not significantly alter tumor growth kinetics or body weight (**Fig. 6B, C and Supplementary Fig. S7D**), indicating limited single-agent activity and good tolerability. IR alone modestly delayed growth in a subset of tumors; however, the overall reduction in tumor volume did not reach statistical significance compared with vehicle-treated controls (**Fig. 6B and C**). In contrast, the combination of AZD7648 and IR produced a pronounced and sustained suppression of tumor growth, significantly exceeding the effects of either treatment alone **(Fig. 6B and C)**. Histopathological analysis supported these findings. H&E staining revealed reduced tumor cellularity in the combination treatment group relative to all other conditions **(Fig. 6D)**. Consistently, quantitative Ki-67 immunostaining demonstrated a significant decrease in proliferative index in tumors treated with AZD7648 plus IR compared with vehicle, AZD7648 alone, or IR alone **(Fig. 6E)**. Collectively, these results demonstrate that DNA-PKcs inhibition with AZD7648 enhances the antitumor efficacy of IR in an IDH1-mutant glioma model *in vivo*. Together with our findings on PARG loss, these findings support a model in which targeting PAR-dependent DNA damage and replication stress pathways can potentiate radiotherapeutic responses in IDH-mutant tumors.

**Figure 6.**
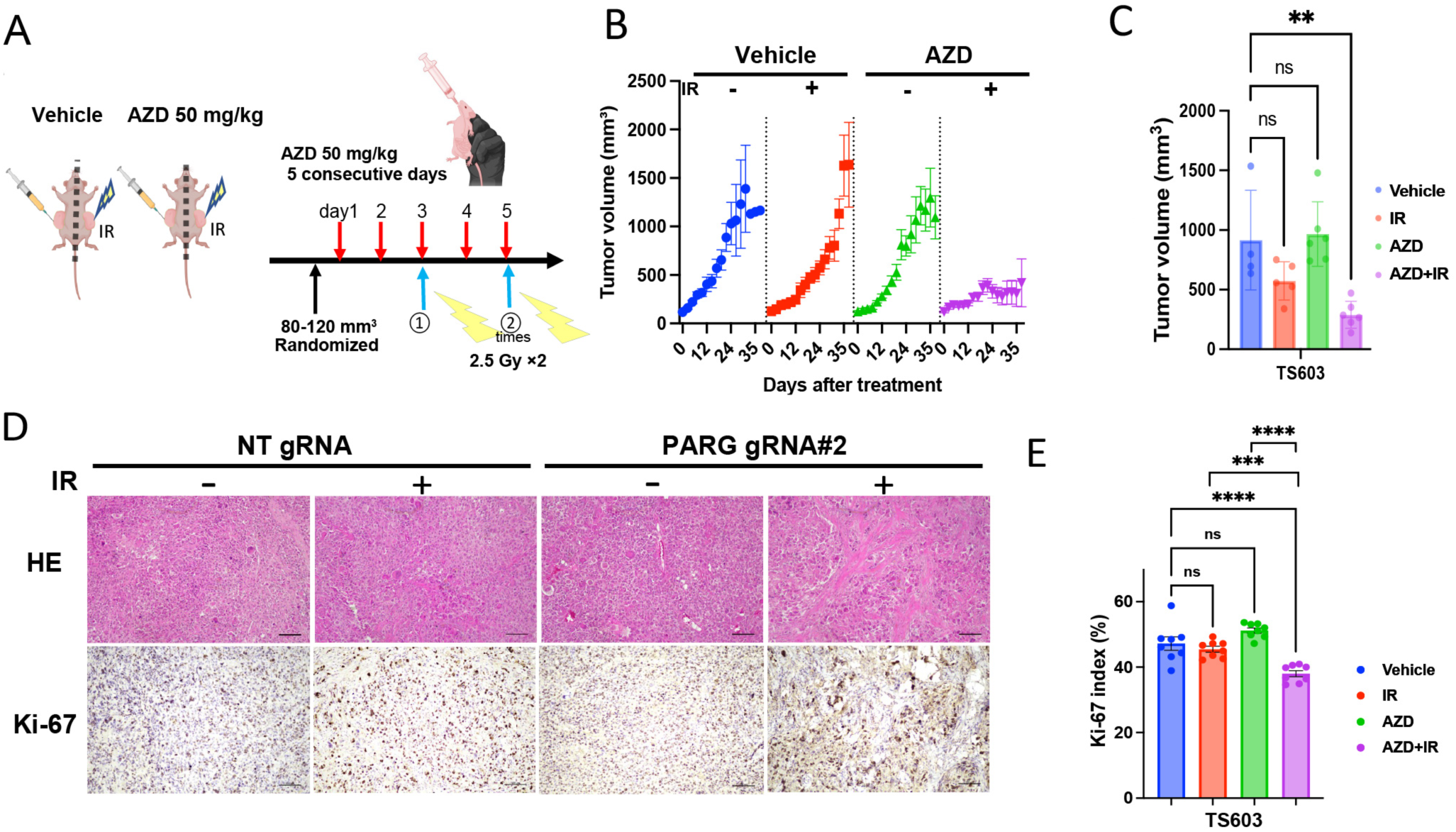

## Discussion

This study establishes that disruption of PAR homeostasis through PARG inhibition or loss selectively amplifies IR cytotoxicity in IDH-mutant glioma, accompanied by perturbed replication tract dynamics and sustained fork-associated signaling through XRCC1 and DNA-PKcs. This enhanced radiosensitivity can be further leveraged by pharmacologic DNA-PKcs inhibition *in vitro* and *in vivo* (**Fig. 7**). These findings build on, and materially extend, the emerging framework in which IDH mutation generates heterochromatin-dependent replication stress without obligating loss of homologous recombination, thereby highlighting PAR-mediated fork stabilization as a relevant S-phase genome maintenance pathway in IDH-mutant cells (18,19). During the S-phase, PARP1 detects unligated Okazaki fragment intermediates and initiates localized PARylation (19). PAR signaling contributes to efficient lagging-strand maturation through XRCC1-dependent repair scaffolds, and timely PAR turnover by PARG supports replication fidelity during S phase (20,26). PARP1 auto-modification has also been implicated in limiting replication fork speed and supporting Okazaki fragment processing, suggesting that defective PAR turnover after PARG loss could perturb lagging-strand processing, chromatin-associated fork dynamics, and replication checkpoint signaling (20,21,27). That these interconnected vulnerabilities are selectively amplified in an IDH-mutant background positions the PARP–PARG axis as a genotype-specific therapeutic node in IDH-mutant glioma, one that is mechanistically distinct from, and complementary to, the alkylation-linked NAD^+^ sequestration strategy we previously described (12). Building on this framework, our results define a PAR homeostasis–linked S-phase vulnerability in IDH-mutant cells in which PARG disruption enhances IR-associated fork signaling and can be further exploited by DNA-PKcs inhibition.

**Figure7.**
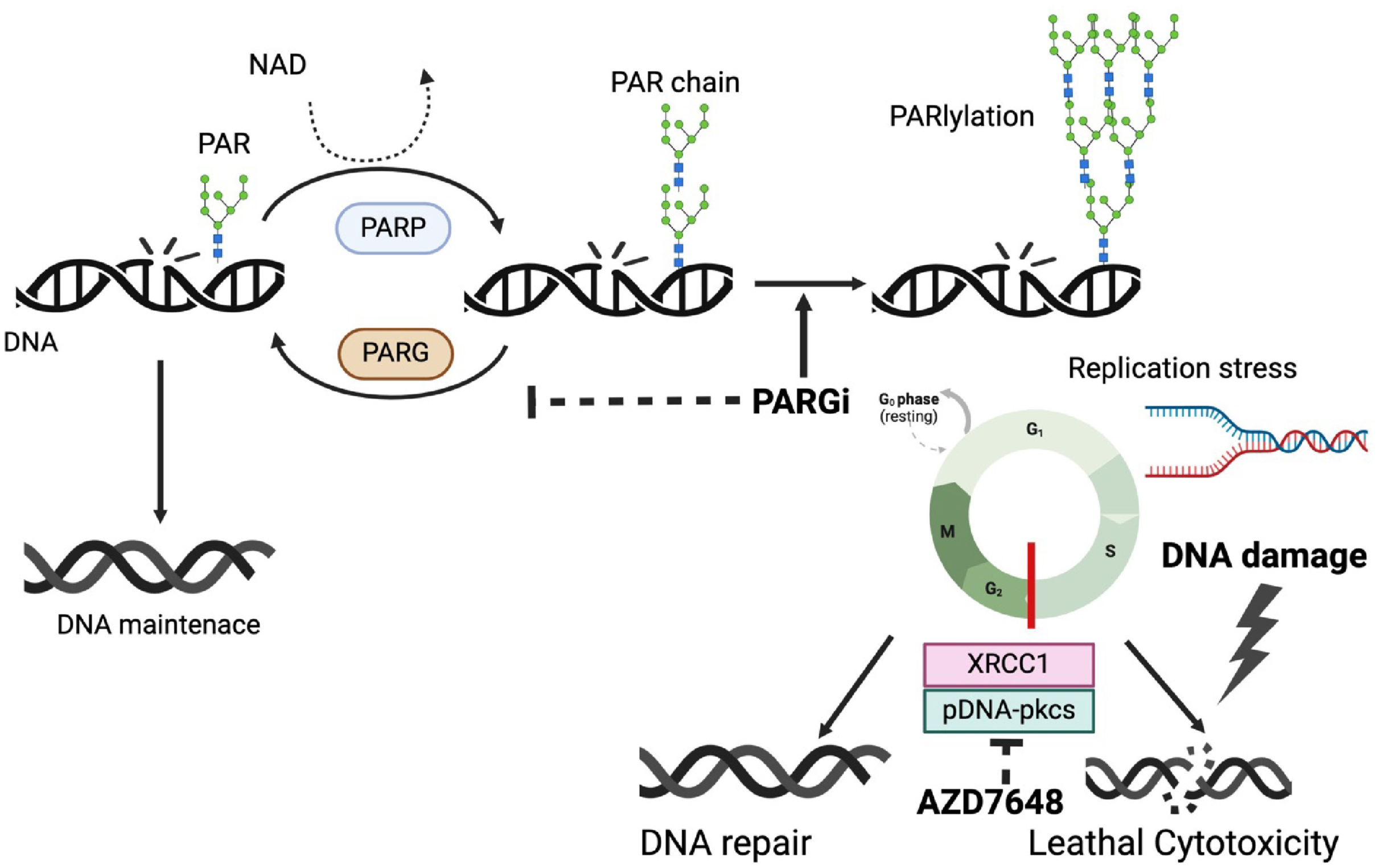

A key mechanistic insight from this study is the formal separation of the radiosensitizing mechanism from the metabolic lethality that underlies alkylation sensitivity. NMN-mediated NAD^+^ repletion, which reverses the cytotoxicity of combined PARG inhibition and TMZ (12), did not rescue the cytotoxicity of combined PARG inhibition and IR (**Fig. 2A–2C**; **Supplementary Fig. S2A–S2C**). This indicated that NAD^+^ depletion alone does not account for the radiosensitizing effect. Rather, these findings separate two therapeutically relevant mechanisms within the same enzymatic axis: alkylation-linked NAD^+^ sequestration and IR-associated replication stress signaling. Consistent with a replication-coupled mechanism, PARG-deficient cells displayed prolonged PAR accumulation after IR, coincident with sustained γH2AX signaling (**Fig. 2D–2H**). Although canonical double-strand break repair, including NHEJ, likely contributes to the IR response, the persistence of γH2AX together with sustained PAR accumulation in PARG-deficient cells points to an additional S-phase-associated, replication-coupled component. In this setting, impaired PAR turnover may delay the resolution of fork-associated repair intermediates at IR-induced template lesions (28,29). Thus, the co-occurrence of sustained PAR and γH2AX after IR supports a model in which PARG disruption amplifies S-phase-associated replication damage signaling in IDH-mutant cells.

DNA fiber analysis demonstrated that PARG loss increased replication tract length relative to non-targeting controls, and that AGI-5198 treatment extended tract length in both backgrounds, with the greatest elongation observed when AGI-5198 was combined with PARG deficiency (**Fig. 3A–3C**). Because PARP1 auto-modification has been reported to limit replication fork speed, the longer replication tracts observed under PARG loss-mediated PAR accumulation appear paradoxical at first glance (21). We propose that this apparent paradox may be resolved by distinguishing instantaneous fork velocity from replication tract length as measured by DNA fiber assay, which captures the total genomic distance replicated per active fork over the labeling period. Under these conditions, individual tracts could appear longer without acceleration of fork velocity, and the observed elongation under PARG deficiency might reflect altered origin usage rather than increased instantaneous speed. An alternative, non-exclusive interpretation is that the PARG-deficient context differs mechanistically from the auto-modification-proficient context. PARP1 is catalytically active but its products accumulate abnormally on chromatin, potentially altering nucleosome compaction and fork-barrier properties in ways not recapitulated by PARP1 auto-PARylation. Distinguishing between these models will require simultaneous quantification of fork speed and inter-origin distance.

Cell-cycle profiling revealed a baseline G1-to-S imbalance in PARG-deficient cells, indicating that impaired PAR turnover perturbs cell-cycle distribution, prior to any exogenous genotoxic stress, consistent with the essential role of dePARylation for the completion of unperturbed S phase (**Fig. 3D–3F**). AGI-5198 more markedly reduced the S-phase fraction in non-targeting cells, whereas IR preferentially depleted the S-phase fraction in PARG-deficient cells. This pattern suggests that PARG disruption alters the balance between replication entry shaped by the 2-HG-associated chromatin state and intra-S-phase stress responses, rendering PARG-deficient cells less able to maintain S-phase progression after IR-induced damagThe differential sen. Correspondingly, IR in PARG-deficient cells induced sustained PAR accumulation alongside elevated XRCC1 and phospho–DNA-PKcs (**Fig. 3G**), consistent with the engagement of the PAR-dependent fork repair scaffold in which PARP1 and DNA-PKcs cooperate to recruit XRCC1 at unresected stalled forks (23). Notably, AGI-5198 attenuated XRCC1 and phospho–DNA-PKcs signals in the PARG-deficient background while simultaneously extending replication tract length. This apparent dissociation may indicate that fork-associated signaling marks primarily the burden of unresolved intermediates at IDH-mutant heterochromatic loci and that mutant IDH inhibition partially relieves this intermediate burden without restoring normal fork-speed dynamics.

The selective elevation of phospho-DNA-PKcs in PARG-deficient cells after IR provided a direct rationale for testing whether DNA-PKcs inhibition could exploit this signaling dependency as a therapeutic strategy. Selective DNA-PKcs inhibition with AZD7648 markedly enhanced IR cytotoxicity in IDH-mutant glioma models *in vitro*, and this radiosensitizing effect translated to effective tumor growth suppression in IDH-mutant xenografts in combination with IR *in vivo* (**Fig. 4A–4E, 6A–6E**; (25)). A mechanistically critical distinction is that DNA-PKcs operates in two separable capacities relevant to this context. As the central kinase of classical NHEJ, DNA-PKcs mediates end-joining of DSBs and is required broadly regardless of cell-cycle phase or replication status (30). As a fork-associated protector, DNA-PKcs promotes replication fork reversal and protects stalled forks from collapse in an NHEJ-independent manner (23,31). The preferential phosphorylation of DNA-PKcs in PARG-deficient cells after IR is most consistent with its recruitment to stalled replication forks, where it cooperates with PARP1 to recruit XRCC1 and coordinate fork-associated repair. AZD7648 treatment would therefore impair both canonical DSB end-joining and this fork-protection function simultaneously, generating a compound repair deficit that IDH-mutant cells are unable to overcome. The differential sensitivity of PARG-deficient, IDH-mutant cells to DNA-PKcs inhibition may therefore reflect combined impairment of canonical DSB repair and fork-protection functions, both of which become more consequential under IR-associated replication stress.

These findings carry substantive clinical relevance for IDH-mutant glioma, where the extended survival characteristic of molecularly favorable subgroups, with median overall survival exceeding a decade in some cohorts (3,32), highlights the importance of both durable tumor control and the minimization of late treatment-related toxicity. The clinical availability of vorasidenib for grade 2 IDH-mutant glioma reshapes this therapeutic context in which PARG-directed radiosensitization would be considered. In related work from our group, short- and long-term mutant IDH1 inhibition did not uniformly induce radioresistance across endogenous patient-derived IDH1-mutant glioma models, although radiation responses varied by genetic context (33). These findings do not support a simple recommendation to withhold mutant IDH inhibition before radiotherapy. Instead, they highlight the need to prospectively evaluate how mutant IDH inhibition, radiotherapy, and PARG-directed radiosensitization should be integrated, with 2-HG suppression, PAR kinetics, and fork-associated signaling serving as candidate pharmacodynamic readouts.

Several important considerations should be noted when interpreting the findings of this study. First, our IR paradigms used subcutaneous xenografts to ensure reproducible tumor measurement and irradiation geometry as an initial proof of concept; however, testing fractionated regimens in orthotopic models will be important to determine the robustness of targeting PAR homeostasis-linked S-phase vulnerabilities. Second, while our data document altered replication tract length and enhanced fork-associated signaling, we did not directly interrogate PARP1 auto-modification status, the distribution of PARP1, or the temporal resolution of PAR accumulation at individual replication forks. Future studies that integrate biochemical assessment of PARP1 modification with orthogonal replication-fork measurements will be required to clarify which replication-control parameters best align with the tract-length phenotype and the sustained XRCC1 and DNA-PKcs signals observed after IR. Third, the context-dependent magnitude of IDH inhibitor effects on replication-stress markers indicates that inhibitor timing, dose, and scheduling may critically influence the PARG-targeting window. Systematic evaluation of inhibitor timing together with pharmacodynamic measures of mutant IDH activity and replication-stress markers is needed.

Finally, translation of PARG-directed radiosensitization requires a clear distinction between target validation and CNS drug delivery. In this study, pharmacologic PARG inhibition was evaluated in vitro, whereas the in vivo radiotherapy experiments relied on genetic PARG loss rather than systemic PARG inhibitor administration. Nevertheless, the therapeutic tractability of PARG inhibition is supported by *in vivo* active preclinical compounds such as COH34, whereas PDD00017273 remains a useful mechanistic tool compound with limitations for systemic *in vivo* application (34). More recently, multiple oral or bioavailable PARG inhibitors have entered early-phase clinical development, including IDE161 (NCT05787587), DAT-2645 (NCT06614751), ETX-19477 (NCT06395519), XNW29016 (NCT06987500), and FORX-428 (NCT07356453), primarily in advanced solid tumors enriched for homologous recombination deficiency, BRCA1/2 alterations, DNA damage response defects, or high replication stress (35–37). However, these programs remain early, and activity in primary CNS tumors, adequate CNS exposure, and compatibility with fractionated radiotherapy have not been established. Therefore, clinical translation to IDH-mutant glioma will require direct assessment of brain exposure, pharmacodynamic PAR suppression, normal brain toxicity, and compatibility with radiotherapy. Similarly, AZD7648 provides a useful mechanistic comparator for DNA-PKcs inhibition in our models, but its first-in-human study reported limited antitumor activity and toxicity concerns in combination with pegylated liposomal doxorubicin (38). In contrast, peposertib provides a more directly brain tumor-relevant clinical precedent because it is being evaluated with radiotherapy in newly diagnosed MGMT-unmethylated glioblastoma, including assessment of blood-brain barrier penetration and pharmacodynamic properties in resected tissue (39).

Together, these findings define PAR homeostasis as a targetable determinant of replication - coupled radiation vulnerability in IDH-mutant glioma and provide a mechanistic foundation for genotype-informed radiosensitization strategies that integrate PARG-directed targeting with DNA-PKcs-mediated repair signaling.

## Methods

### Cell Lines and Culture Conditions

Patient-derived glioma sphere lines MGG18, MGG119, and MGG152 were established at Massachusetts General Hospital from surgical specimens obtained under IRB-approved protocols (40,41). MGG18 cells stably expressing tetracycline-inducible *IDH1*-R132H (MGG18-*IDH1*-R132H) were generated by lentiviral transduction as previously described (11).

TS603, a WHO grade III *IDH1*-R132H-mutant oligodendroglioma line with 1p/19q co-deletion, was obtained as previously described (42). All experiments labeled as TS603 in this study used TS603S2, a subclone of TS603 selected for stable IDH1 R132H expression; for simplicity, this line is referred to as TS603 throughout the manuscript(12,33). HT1080, an *IDH1*-R132C-mutant fibrosarcoma line, was obtained from ATCC. MGG18, MGG119, MGG152, and TS603 were cultured in Neurobasal medium supplemented with B27, EGF (20 ng/mL), FGF (20 ng/mL), heparin, GlutaMAX, and penicillin–streptomycin. HT1080 cells were cultured in EMEM supplemented with 10% FBS and penicillin–streptomycin. All cells were maintained at 37 °C in a humidified atmosphere containing 5% CO₂. Cell lines were confirmed to be mycoplasma-free by PCR testing (2020–2023). HT1080 cells were authenticated by short tandem repeat (STR) profiling (2023).

### Compounds and Chemicals

The following compounds were used: AGI-5198 (Cayman Chemical), AZD7648 (HY-111783; MedChemExpress), dimethyl sulfoxide (DMSO; Sigma-Aldrich), doxycycline hyclate (Sigma-Aldrich), lomustine (CCNU; Sigma-Aldrich), nicotinamide (Sigma-Aldrich), β-nicotinamide mononucleotide (NMN; Sigma-Aldrich), PDD00017273 (PDD; Tocris Bioscience), and temozolomide (TMZ; Sigma-Aldrich).

### CRISPR/Cas9-Mediated PARG Knockout

Guide RNAs targeting *PARG* (gRNA #2: 5ʹ-TGCTATTCTGAAATACAATG-3ʹ; gRNA #3: 5ʹ-CAACACATTATAAAGATTTG-3ʹ) and a non-targeting control (NT: 5ʹ-ATCTACGGGTTTATGCCAAT-3ʹ) were cloned into pLentiCRISPR v2. Lentivirus was produced in HEK293T cells. HT1080 cells were transduced in the presence of polybrene (8 µg/mL) and selected with puromycin (0.6 µg/mL). PARG knockout was confirmed by Western blot.

### Irradiation

Cells and tumor-bearing mice were irradiated with ionizing radiation (IR) using an X-RAD 320 X-ray irradiator (Precision X-Ray, North Branford, CT) operated at 320 kVp and 12.5 mA with a 2.0 mm aluminum filter at a source-to-sample distance of 50 cm. The dose rate was approximately 2.3–2.4 Gy/min. Cells were irradiated at room temperature in white 96-well plates containing 100 µL medium per well, or in culture dishes as specified for each assay. For *in vivo* experiments, a single dose of 3 Gy (HT1080 model) or a total of 5 Gy delivered in two fractions (TS603 model) was administered to the right flank.

### Cell Viability Assay

Cells were seeded at 1–3 × 10³ cells per well in 96-well plates, treated with the indicated compounds, irradiated, and cultured for 72–120 h. Viability was measured using the CellTiter-Glo Luminescent Cell Viability Assay (Promega) according to the manufacturer’s instructions at the indicated time points.

### Clonogenic Assay

HT1080 cells were seeded at 100 cells per well in 6-well plates. Cells were exposed to PDD (5 µM) or DMSO for 18 h, irradiated at the indicated doses, and cultured for 7–14 days. Cells were fixed with 6% glutaraldehyde and stained with 0.5% crystal violet. Images were captured with the EVOS™ FL Auto 2 Imaging System (Thermo Fisher Scientific).

### Limiting Dilution Assay

MGG18 cells were seeded at 1–20 cells per well in 96-well plates, treated with PDD (5 µM) or DMSO, irradiated, and cultured under spheroid conditions for 7 days. Sphere formation was scored, and stem-cell frequency was calculated using ELDA software (http://bioinf.wehi.edu.au/software/elda/).

### NAD^+^ Quantitation

Intracellular NAD^+^ levels were measured using the NAD/NADH-Glo Assay (Promega). Briefly, 1 × 10⁵ cells were lysed, and lysates were processed for selective NAD^+^ or NADH detection. Luminescence was measured in 96-well plates. Detailed lysis and detection procedures are provided in Supplementary Methods.

### Comet Assay

HT1080 cells were treated with 5 Gy IR or TMZ, with 100 µM tert-butyl hydroperoxide (tBH) as a positive control. Alkaline comet assays were performed using the CometAssay Reagent Kit (Trevigen, 4250-050-K) according to the manufacturer’s instructions. Cells were embedded in low-melting-point agarose on CometSlides, lysed, subjected to alkaline unwinding (20 min), and electrophoresed in Alkaline Electrophoresis Solution at 1 V/cm for 30 min at 4 °C. After washing and drying, samples were stained with SYBR Green I. Comet images were acquired by epifluorescence microscopy and analyzed using the OpenComet plugin for ImageJ (43). Detailed electrophoresis and staining conditions are provided in Supplementary Methods.

### Western Blot

Cells were lysed in RIPA buffer, and 10 µg of total protein per lane was resolved on 4–20% SDS-PAGE gels and transferred to PVDF membranes. Membranes were probed with the following primary antibodies: anti-PAR, anti-PARG, anti-γH2AX, anti-β-actin, anti-IDH1-R132H, anti-XRCC1, anti-Cyclophilin B, anti-phospho-DNA-PKcs, anti-cleaved caspase-3, and anti-Vinculin. Catalog numbers, dilutions, and detailed protocols are provided in Supplementary Methods.

### EdU Cell-Cycle Analysis

HT1080 cells were pulse-labeled with 5-ethynyl-2’-deoxyuridine (EdU), irradiated with 5 Gy IR, and processed using the Click-iT EdU Alexa Fluor 647 Flow Cytometry Assay Kit (Thermo Fisher Scientific). Cells were counterstained with propidium iodide (PI)/RNase buffer and analyzed on a BD LSR II flow cytometer. Data were analyzed with FlowJo (version 10.10.0; BD Life Sciences); S-phase fraction was quantified as the percentage of EdU-positive cells. Detailed procedures are provided in Supplementary Methods.

### DNA Fiber Assay

HT1080 cells (0.5 × 10⁶) were seeded in 6-well plates, sequentially pulse-labeled with chlorodeoxyuridine (CldU; 100 µM, 20 min) and iododeoxyuridine (IdU; 250 µM, 30 min), harvested, and resuspended in chilled PBS (∼2,500 cells/µL). Cell suspensions (3 µL) were lysed on glass slides with fiber lysis buffer (200 mM Tris-HCl pH 7.4, 50 mM EDTA, 0.5% SDS), and DNA fibers were spread by gravity. Slides were fixed in methanol/acetic acid (3:1), denatured in 2.5 N HCl for 1 h, blocked in 2% BSA/0.1% Tween 20 in PBS, and immunostained with rat anti-BrdU (ab6326, Abcam; 1:100, detecting CldU) and mouse anti-BrdU (347580, BD Biosciences; 1:50, detecting IdU), followed by donkey anti-rat Alexa Fluor 594 and donkey anti-mouse Alexa Fluor 488 secondary antibodies (1:100). Slides were mounted with ProLong Gold Antifade (Thermo Fisher Scientific). Fiber images were acquired by fluorescence microscopy, and total tract lengths (CldU + IdU) were measured using ImageJ. A minimum of 291 fibers per condition were analyzed from three independent experiments (total 1,529 fibers across four conditions).

### Immunofluorescence Staining

HT1080 cells (NT gRNA, PARG gRNA #2, and PARG gRNA #3) were plated on poly-D-lysine– coated glass coverslips in 24-well plates (∼76,000 cells per well) in EMEM supplemented with 10% FBS. Cells were treated with or without IR and fixed with 4% paraformaldehyde in PBS. After PBS washes, cells were permeabilized and blocked with 5% BSA/0.3% Triton X-100 in PBS for 1 h at room temperature. Cells were then incubated overnight at 4 °C with primary antibodies against PAR and 19 hosphor-Histone H2A.X (Ser139) diluted in blocking buffer. The following day, after PBS washes, cells were incubated with Alexa Fluor 488– and Alexa Fluor 594– conjugated secondary antibodies for 2 h at room temperature. Images were acquired by fluorescence microscopy, and fluorescence intensity was quantified using ImageJ. Data are presented as mean ± SEM from three independent experiments (n = 3 per condition per time point). Antibody details are listed in Supplementary Table S1; dilution and detailed protocols are provided in Supplementary Methods.

### Immunohistochemistry

Formalin-fixed, paraffin-embedded tumor sections were stained with hematoxylin and eosin (H&E) for morphological assessment or with anti-Ki-67 antibody (1:200) followed by HRP-based detection. For Ki-67 quantification, six images per section were captured at 20× magnification, and the percentage of Ki-67–positive nuclei was calculated.

### In Vivo Studies

All animal experiments were approved by the Institutional Animal Care and Use Committee (IACUC) of Massachusetts General Hospital. Female athymic nude mice (7–10 weeks old, 21– 26 g) were used.

### HT1080 and TS603 bilateral flank model

NT (2 × 10⁶) and PARG KO (4 × 10⁶) HT1080 cells were resuspended in 200 µL of EMEM/Matrigel (1:1) and implanted subcutaneously into both flanks of the same mouse (n = 10 mice). Once tumors reached approximately 120 mm³, one flank tumor per mouse received focal IR, while the contralateral tumor served as the non-irradiated control.

All experiments labeled as TS603 in this study used TS603S2, a subclone of TS603 selected for stable IDH1 R132H expression; for simplicity, this line is referred to as TS603 throughout the manuscript (12,33). TS603 cells (2 × 10⁶) were resuspended in 200 µL of DMEM/Matrigel (1:1) and implanted into both flanks (n = 12 mice). Once tumors reached approximately 80-120 mm³,

AZD7648 (50 mg/kg) was administered to the treatment groups once daily by oral gavage for five consecutive days. Then, one flank tumor per mouse received focal IR, while the contralateral tumor served as the non-irradiated control. Tumor and body weight measurements were performed according to our previous report (12).

### Statistical Analysis

Data are presented as mean ± SEM unless otherwise stated. Two-group comparisons were performed using a two-sided Student’s t-test. Multi-group comparisons were analyzed by one-way or two-way ANOVA followed by Tukey’s, Dunnett’s, or Šídák’s multiple comparisons test, as indicated in each figure legend. DNA fiber tract-length data were analyzed by Welch’s ANOVA followed by Dunnett’s T3 multiple comparisons test. Comet assay tail moments were analyzed by two-way ANOVA followed by Šídák’s multiple comparisons test. Immunofluorescence quantification data were analyzed by two-way ANOVA followed by Tukey’s multiple comparisons test. Limiting-dilution assay data were analyzed using ELDA software (http://bioinf.wehi.edu.au/software/elda/). All statistical analyses were performed using GraphPad Prism (version 11.0.0; RRID:SCR_002798) (v11). A p-value < 0.05 was considered statistically significant.

### Data Availability

All data generated in this study are available within the article and its supplementary data files. Raw data are available from the corresponding authors upon reasonable request.

## Authors’ Contributions

Conception and design: YK, HW, DPC

Development of methodology: YK, AN, EW, AK, LM, CCC, HN, JJM

Acquisition of data (provided animals, acquired and managed patients, provided facilities, etc.): YK, AN, EW, AK, LM, CCC

Analysis and interpretation of data (e.g., statistical analysis, biostatistics, computational analysis): YK, AK

Writing, review, and/or revision of the manuscript: YK, AN, AK, HW, DPC Study supervision: JJM, HW, DPC

## Acknowledgments

We would like to thank Kensuke Tateishi for establishing the MGG18 cell line with the tetracycline-inducible system, and Logan Muzyka for assisting with the processing of samples for the animal experiments

