## Supplementary figures and images for "Augmenting Radiation Sensitivity by Targeting PAR-Dependent Replication Fork Vulnerability in *IDH-*Mutant Glioma"

### Supplemental FigS1

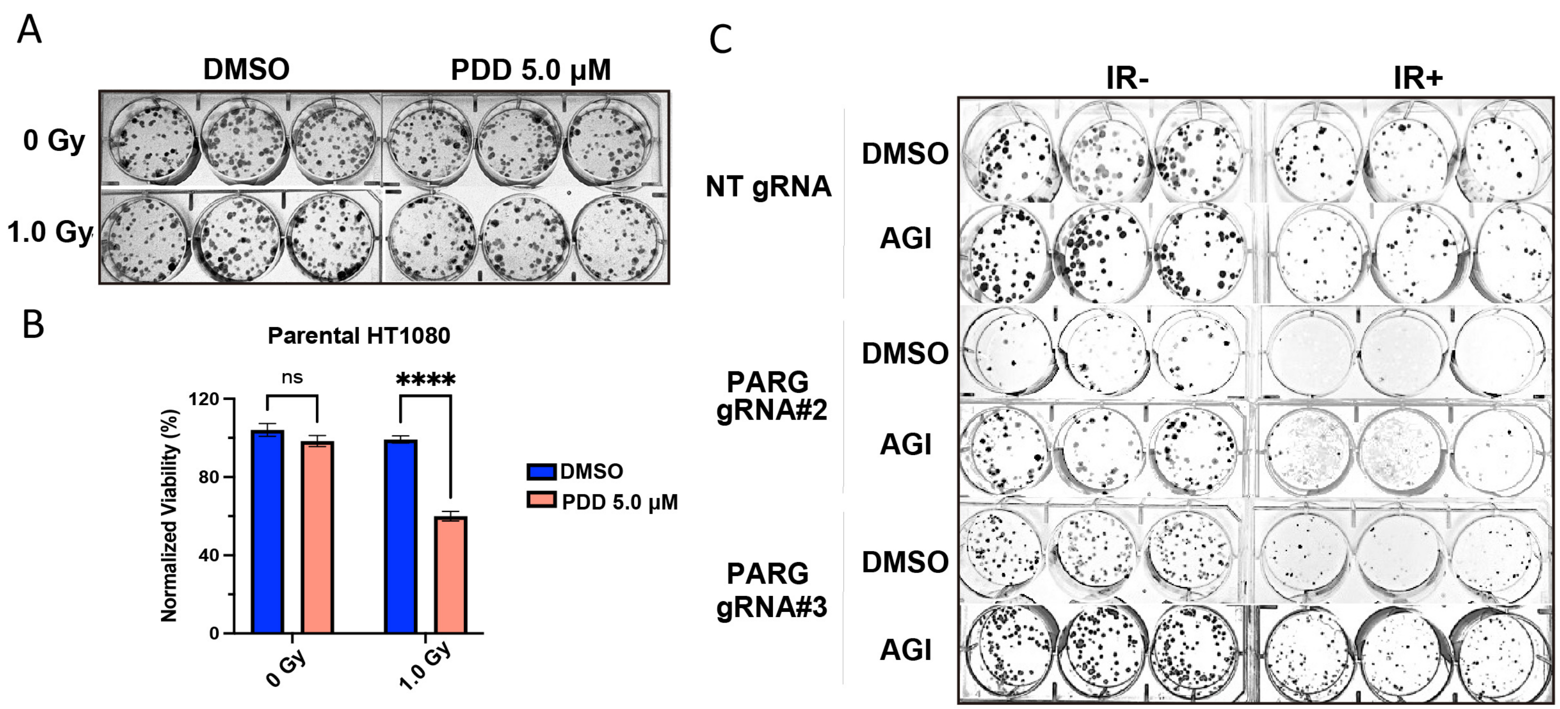

### Supplemental FigS2

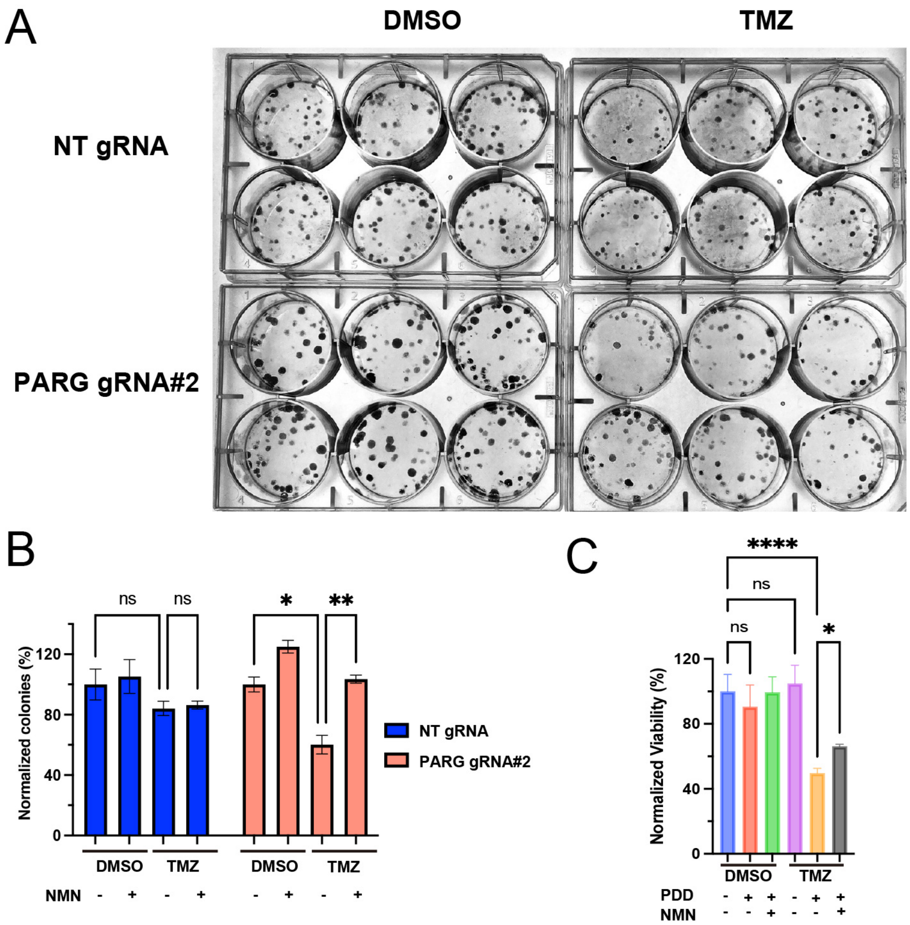

### Supplemental FigS3

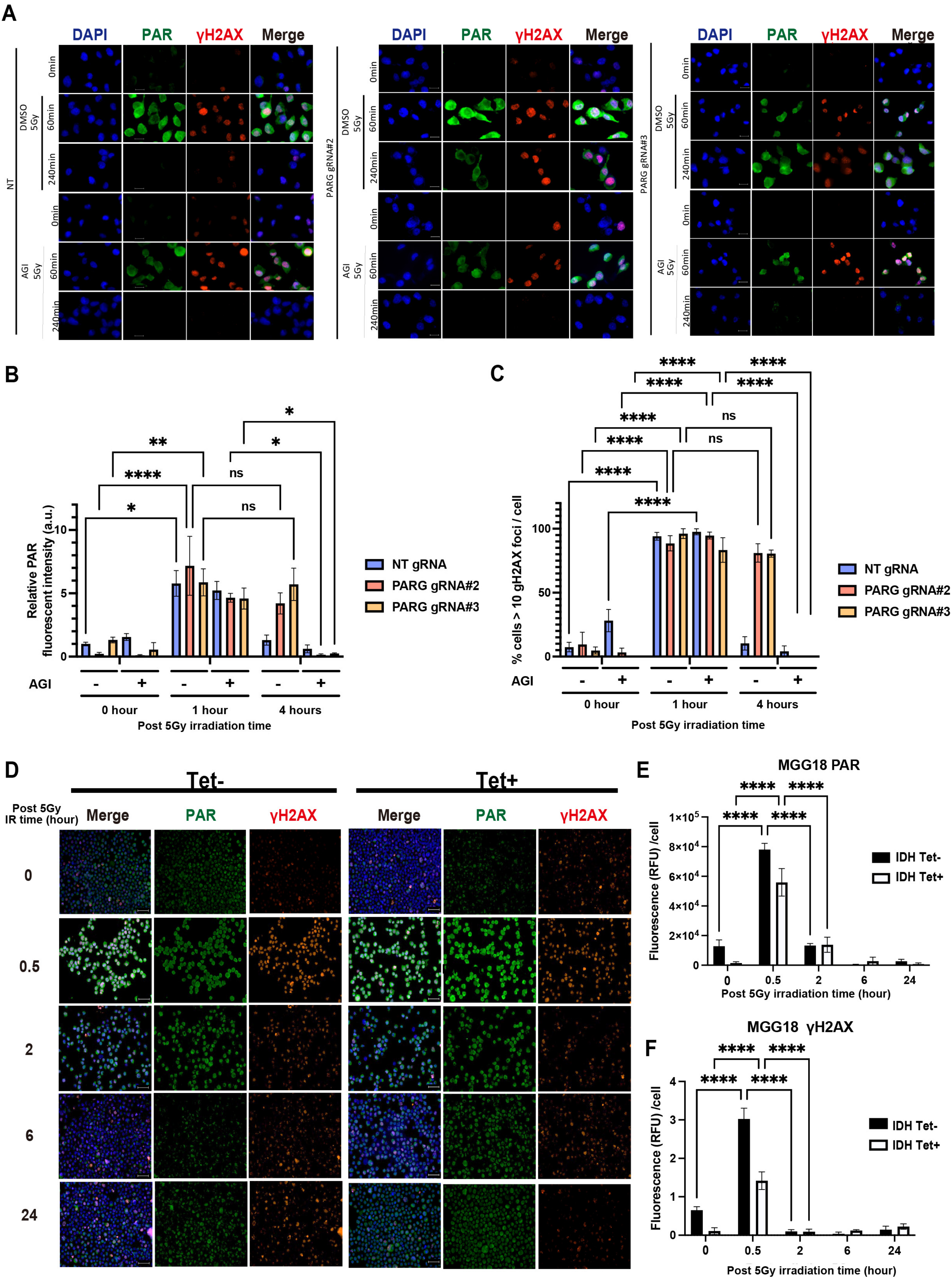

### Supplemental FigS4

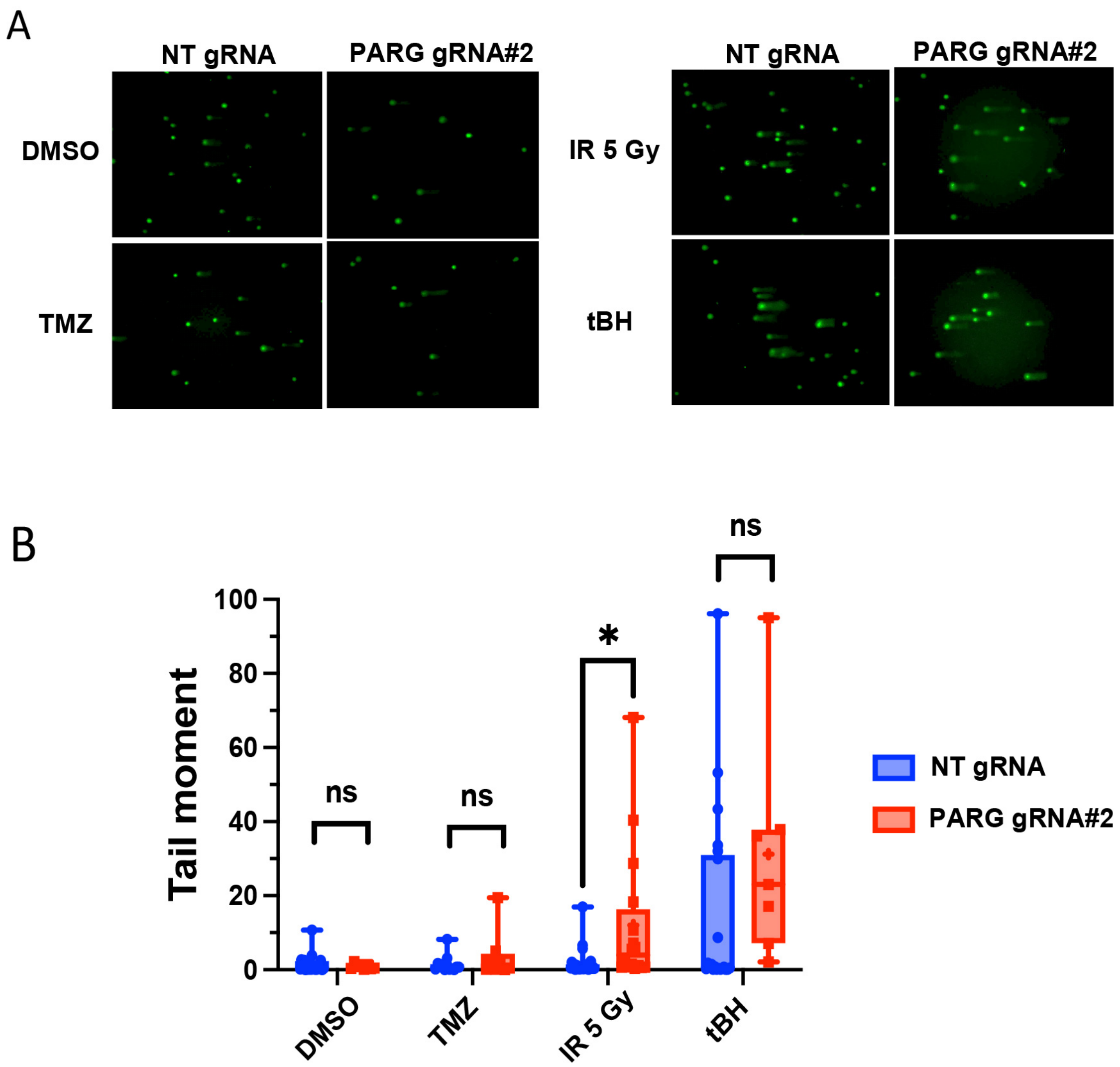

### Supplemental FigS5

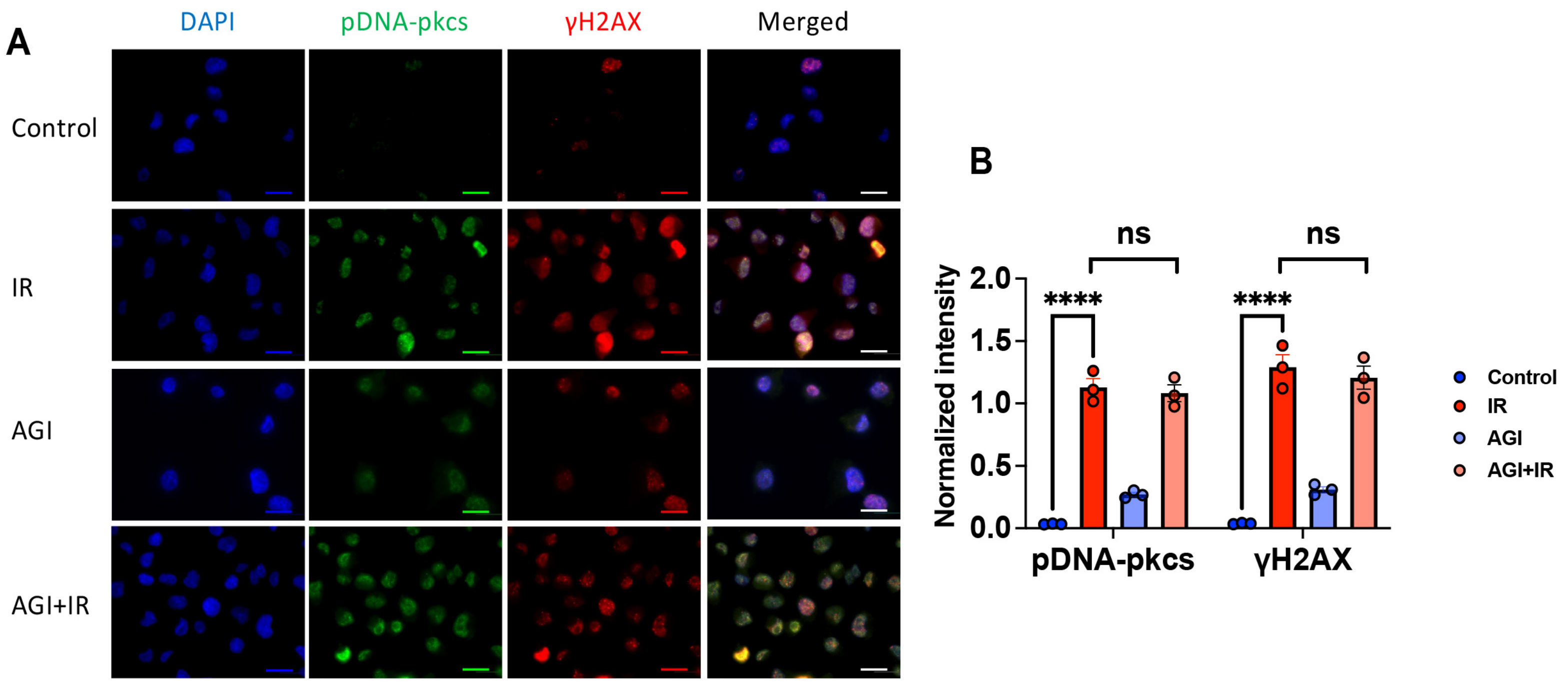

### Supplemental FigS6

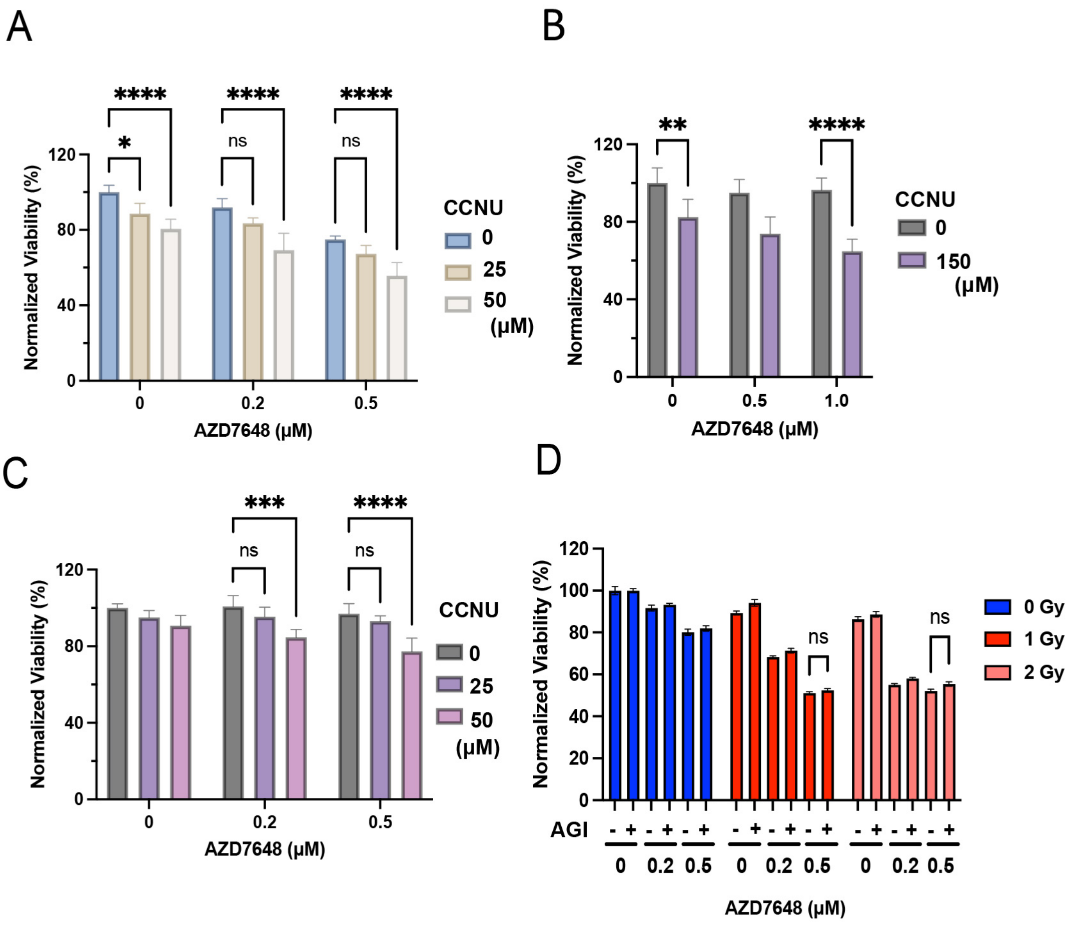

### Supplemental FigS7

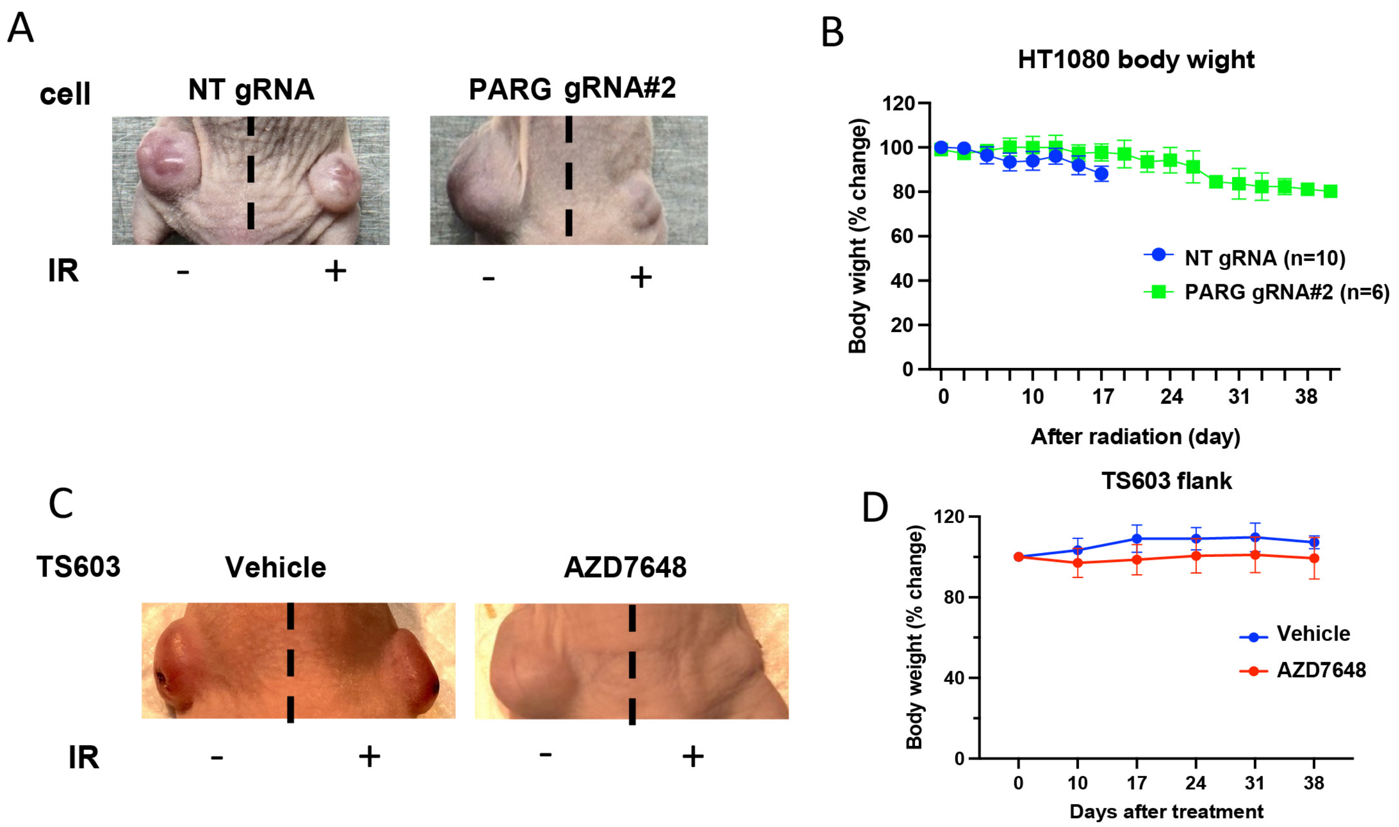
