## Supplemental information for "Augmenting Radiation Sensitivity by Targeting PAR-Dependent Replication Fork Vulnerability in *IDH-*Mutant Glioma"

**Supplementary figure S1: Pharmacologic and genetic PARG inhibition enhances radiation-induced clonogenic cell death in HT1080 cells**.

(**A**) Clonogenic assay of parental HT1080 after 7 days exposure to either 0 or 1 Gy of ionizing radiation (IR) with DMSO or 5.0 μM PDD00017273 (PDD). Representative images of single-cell clone proliferation, stained with crystal violet. (**B**) Quantification of colony number from conditions represented in panel (**A**). (**C**) Clonogenic assay of NT, PARG gRNA#2, and PARG gRNA#3 HT1080 cells treated with or without AGI-5198 (AGI) (5 μM) and exposed to 0 or 1.5 Gy IR. Representative crystal-violet-stained colony images. Statistical analyses were performed using Unpaired two-tailed Student's *t-test*. Data are presented as mean ± SEM. Statistical significance indicated as ns (not significant), ****$p<0.0001$.

**Supplementary figure S2: $NAD^+$ repletion by NMN partially rescues TMZ-induced clonogenic cell death in PARG-deficient HT1080 cells.**

(**A**) Clonogenic assay of NT and PARG knockout (KO) HT1080 cells after 7 days exposure to Temozolomide (TMZ) (200 $\mu$M) with or without Nicotinamide Mononucleotide (NMN) (1 mM). Representative images depict single-cell clone proliferation, stained with crystal violet. (**B**) Quantification of colony number from various conditions represented in (A), normalized to the number of colonies in the control condition of DMSO and without NMN. (**C**) CellTiter-Glo assay (CTGA) of parental HT1080 cells after 72 hours of exposure to TMZ with or without 5.0 μM PDD00017273 (PDD) and NMN. Statistical analyses were performed using two-way ANOVA. Data are presented as mean ± SEM. Statistical significance indicated as ns (not significant), *$p<0.05$, **$p<0.01$, ****$p<0.0001$.

**Supplementary figure S3: Comparable PAR and γH2AX dynamics following irradiation in IDH-wildtype and IDH-mutant MGG18 cells.**

(**A**) Immunofluorescence microscopy images depicting PAR and γ-H2AX foci formation of HT1080 NT gRNA, PARG gRNA #2, and #3 cells before and after exposure to 5 Gy irradiation (IR). Cells were treated with IR ± IDH inhibitor AGI-5198 (5 μM). Confocal microscopy was used to detect PAR (green), γ-H2AX foci (red), and nuclei stained with DAPI (blue). Scale bar: 20 μm. (**B**) Quantification of average fluorescence intensity of PAR from (A). (**C**) Percentage of cells displaying more than 10 γ-H2AX foci per cell at various time points post 5 Gy IR from (C). The mean and standard error of the mean from three independent experiments are displayed. Units are in arbitrary units (a.u.). (**D**) Immunofluorescence analysis of PAR (Poly ADP-ribose) and γH2AX (phosphorylated histone H2AX) levels in MGG18s cells with regulated IDH1R132H expression using the tetracycline-inducible (tet) system. The captured images at different time points after 5 Gy irradiation (0h, 0.5h, 2h, 6h, and 24h) illustrate the staining patterns of PAR, γH2AX, and their merge for MGG18 tet(-) and MGG18 tet(+) cells. Scale bar: 50 μm. (**E**) Quantification of PAR levels from the data presented in (A). The X-axis represents the time frames, and the Y-axis represents the fluorescence intensity (RFU, Relative Fluorescence Units) per cell in PAR. (**F**) Quantification of γH2AX foci from the data presented in (A). The X-axis represents the time frames, and the Y-axis represents the number of γH2AX foci per cell. Statistical analyses were performed using two-way ANOVA. Data are presented as mean ± SEM. Statistical significance indicated as ****$p<0.0001$.

**Supplementary figure S4: PARG loss enhances and prolongs irradiation-induced DNA damage in IDH-mutant cells.**

(**A**) Comet assay fluorescent images captured at 4X magnification, comparing the effects of different treatments on DNA damage. NT gRNA and PARG KO HT1080 cells were exposed to DMSO (control), TMZ (200μM) for 24hrs, irradiation, and 100 μM tert-Butyl hydroperoxide (tBH) for 3hrs serving as positive control. (**B**) Quantification of DNA damage observed in (A). The Y-axis represents the tail moment, indicating the extent of DNA damage. Data are presented as mean ± SEM. Statistical analyses were performed using one-way ANOVA. Statistical significance indicated as ns (not significant), *$p<0.05$, **$p<0.01$, ***$p<0.001$, ****$p<0.0001$.

**Supplementary Fig. S5. PARG deficiency accelerates replication fork progression but promotes aberrant and sustained DNA Damage signaling following IR**

(**A**) Immunofluorescence analysis of DNA damage markers in parental HT1080 cells, showing DAPI staining, pDNA-PKcs activation, and γH2AX expression and co-localization in IR only, AGI-5198 (AGI) treatment, and combined AGI+IR conditions. Scale bar: 20 µm. (**B**) Quantification of average fluorescence intensity of pDNA-PKcs and γH2AX in (**A**). Statistical analyses were performed using two-way ANOVA. Data are presented as mean ± SEM. Statistical significance indicated as ns (not significant), *$p<0.05$, **$p<0.01$, ***$p<0.001$, ****$p<0.0001$.

**Supplementary figure S6: DNA-PKcs inhibition confers limited sensitization to CCNU in IDH-mutant glioma cells.**

(**A-C**) These panels present the cell viability analysis using CellTiter-Glo assay (CTGA) in TS603 (**A**), MGG119 (**B**), and MGG152 (**C**) cells after 120-hour exposure to a combination of AZD7648 (AZD) and CCNU. AZD was used at concentrations between 0 to 1.0 µM, while CCNU was used at concentrations between 0 to 150 µM. (**D**) CTGA of combined treatment with AZD and irradiation, alongside AGI-5198 (AGI), in TS603 cells. Data are presented as mean ± SEM. Statistical analyses were performed using one-way ANOVA. Statistical significance indicated as ns (not significant), *$p<0.05$, **$p<0.01$, ***$p<0.001$, ****$p<0.0001$.

**Supplementary figure S7: *In vivo* tumor images and body weight monitoring in PARG-deficient and DNA-PKcs–inhibited xenograft models.**

(**A**) Representative images of tumors transduced with non-targeting (NT) gRNA and PARP gRNA#2, with or without irradiation. Each mouse displays a control (left side, no IR) and treated (right side, IR) condition for their respective tumors. (**B**) Body weight of mice in non-targeting (NT) and PARG knockout PARG gRNA #2 HT1080 flank xenograft groups. (**C**) Representative images of a TS603-S2 flank tumor model comparing the effects of vehicle (left mouse) and AZD-7648 treatment (right mouse). Each mouse displays a control (left side, no IR) and treated (right side, IR) condition for their respective tumors. (**D**) Body weight of mice in vehicle and AZD-7648 (AZD) treated TS603-S2 flank tumor model groups.

### Supplementary Methods

#### Western Blot (Detailed Protocol)

Cells were lysed in RIPA buffer (Thermo Fisher Scientific) supplemented with protease and phosphatase inhibitor cocktails (Roche). Protein concentration was determined by BCA assay. Ten micrograms of total protein per lane were resolved on 4–20% Mini-PROTEAN TGX precast gels (Bio-Rad) and transferred to PVDF membranes. Membranes were blocked in 5% non-fat milk or 5% BSA in TBS-T (0.1% Tween-20), as appropriate for each primary antibody (see Supplementary Table S1), for 1 h at room temperature and incubated overnight at 4 °C with primary antibodies listed in Supplementary Table S1. After washing, membranes were incubated with HRP-conjugated secondary antibodies (1:5,000) for 1 h at room temperature. Bands were visualized using ECL substrate (Bio-Rad) on a Gel Doc XRS+ imaging system (Bio-Rad, RRID:SCR_019690). Images were acquired with Image Lab software (Bio-Rad, RRID:SCR_014210).

#### Immunofluorescence Staining (Detailed Protocol)

HT1080 cells (NT gRNA, PARG gRNA #2, and PARG gRNA #3) were plated on poly-D-lysine–coated round glass coverslips (Fisherbrand) in 24-well plates at a density of ~76,000 cells per well in EMEM with 10% FBS. Following treatment with or without IR, cells were fixed at the indicated time points with 4% paraformaldehyde in PBS for 20 min at room temperature. After three PBS washes, cells were permeabilized and blocked with 5% BSA/0.3% Triton X-100 in PBS for 1 h at room temperature. Cells were incubated overnight at 4 °C with the following primary antibodies diluted in blocking buffer: mouse monoclonal anti-PAR (clone 10H; Enzo Life Sciences, ALX-804-220-R100; 1:100) and rabbit polyclonal anti-phospho-Histone H2A.X (Ser139) (Cell Signaling Technology, #2577; 1:100). The following day, coverslips were washed three times with PBS and incubated for 2 h at room temperature with donkey anti-mouse IgG Alexa Fluor 488 (Invitrogen, A-21202; 1:1,000) and donkey anti-rabbit IgG Alexa Fluor 594 (Invitrogen, A-21207;

1:1,000). PAR was visualized in the 488 nm channel (green) and γH2AX in the 594 nm channel (red). Coverslips were mounted with VectaShield mounting medium containing DAPI (Vector Laboratories, H-1200). Images were captured on a Photometrics fluorescence microscope; a minimum of three random high-power fields were acquired per coverslip. Fluorescence intensity was quantified using ImageJ (NIH; RRID:SCR_003070). Data are shown as mean ± SEM from three independent experiments (n = 3 per condition per time point).

**EdU/PI Flow Cytometry (Detailed Protocol)**

HT1080 cells were pulse-labeled with EdU (15 μM) and irradiated with 5 Gy. Cells were harvested, fixed, and permeabilized using the Click-iT EdU Alexa Fluor 647 Flow Cytometry Assay Kit (Thermo Fisher Scientific, C10424) according to the manufacturer's instructions. EdU was detected by Click-iT reaction with Alexa Fluor 647 azide. Cells were then stained with PI/RNase buffer for DNA content analysis. Samples were acquired on a BD LSR II flow cytometer (RRID:SCR_002159). Single cells were gated based on FSC-A/FSC-H, and a minimum of 10,000 events per sample were collected. Data were analyzed with FlowJo (version 10.10.0; BD Life Sciences; RRID:SCR_008520) using a predefined single-cell gating strategy; S-phase fraction was defined as the percentage of EdU-positive cells.

**$NAD^+$ Quantitation (Detailed Protocol)**

Intracellular $NAD^+$ levels were measured using the NAD/NADH-Glo Assay (Promega). Cells ($1 \times 10^5$) were lysed in 100 μL of PBS containing 1% DTAB (dodecyltrimethylammonium bromide). For selective $NAD^+$ detection, 50 μL of lysate was treated with 25 μL of 0.4 N HCl and heated at 60 °C for 15 min. For NADH detection, 50 μL of lysate was treated with 25 μL of 0.5 M Trizma base and heated at 60 °C for 15 min. After neutralization, samples were transferred to a white 96-well plate and mixed with an equal volume of NAD/NADH-Glo Detection Reagent. Luminescence was measured after 30 min of incubation at room temperature.

**Comet Assay (Detailed Protocol)**

Alkaline comet assays were performed using the CometAssay Reagent Kit (Trevigen, 4250-050-K). LMAgarose was melted in boiling water for 5 min and cooled to 37 °C in a water bath for at least 20 min. HT1080 cells ($1 \times 10^5$/mL in $Ca^{2+}$/$Mg^{2+}$-free PBS) were combined with molten LMAgarose at a 1:10 (v/v) ratio, and 50 μL was pipetted onto each CometSlide sample area.

Slides were placed flat at 4 °C in the dark for 10 min to allow gelling, then immersed in pre-chilled Lysis Solution at 4 °C for 30–60 min. Slides were transferred to freshly prepared Alkaline Unwinding Solution (200 mM NaOH, 1 mM EDTA, pH >13) for 20 min at room temperature in the dark. After draining, slides were placed in the CometAssay ES II electrophoresis tank containing 850 mL of freshly prepared, pre-chilled (4 °C) Alkaline Electrophoresis Solution (200 mM NaOH, 1 mM EDTA, pH >13). Electrophoresis was performed at 1 V/cm (21 V) for 30 min. Slides were then washed twice in deionized water for 5 min each, followed by 70% ethanol for 5 min, and dried at 37 °C for 10–15 min. Dried samples were stained with diluted SYBR Green I (100 µL per sample area) for 5 min at 4 °C, and excess stain was removed by gentle tapping. Slides were air-dried in the dark and imaged by epifluorescence microscopy (excitation 494 nm / emission 521 nm). Tail moments were quantified using the OpenComet plugin for ImageJ (43). Tail moments were quantified from a total of 122 comets across four treatment conditions and two genotypes in three independent experiments.
